# 3D RNA profile of habenulae in the catshark *Scyliorhinus canicula*: a reference in jawed vertebrates

**DOI:** 10.1101/2023.10.17.562684

**Authors:** L. Michel, H. Mayeur, A Quillien, N. Heier, J. Clairet, S. Dejean, P. Blader, R. Lagadec, S. Mazan

## Abstract

We report the generation of a spatially resolved RNA profile of the habenulae in a chondrichthyan, the catshark *Scyliorhinus canicula*. The digital profiles obtained for candidate markers of the broad territories previously characterized in this species reflect their broad characteristics, including left- or right asymmetric distributions. We use spatial autocorrelation and autocorrelation as measures to analyze these data and identify novel territory markers. These analyses lead to a refined molecular characterization of catshark habenulae and lead to extended lists of markers for the major territories previously identified. They confirm the marked asymmetry of the catshark habenulae, both in its lateral and medial components. In addition, they highlight a bilateral lateral to medial organization in radial bands, which may reflect a temporal regulation of neurogenesis. These data lead to a novel reference to explore the lmode of evolution of habenulae and of their asymmetries across gnathostomes.

## 1. INTRODUCTION

The habenulae are bilateral epithalamic structures, present in all vertebrates. They are part of a highly conserved brain circuitry, integrating sensory information and corticolimbic inputs to regulate a variety of behaviors, cognitive and emotional processes, including responses to threats, social behaviors, exercise motivation, reward or aversive learning (Reviewed in (Hu, Cui and Yang, 2020); Okamoto et al., 2021; (Roman *et al.*, 2020)). The complexity of these functions is reflected by a complex subdomain organization, which has thus far been detailed in only two species, the mouse and the zebrafish. In the former, habenulae are partitioned into two broad subdivisions, a medial one (MHb) and a lateral one (LHb), which extensively differ by their cytoarchitecture, molecular properties and projection pattern, respectively to the interpeduncular nuclei and the dorsal raphe (Aizawa *et al.*, 2012; Wagner *et al.*, 2016; Molas *et al.*, 2017; Zhou *et al.*, 2017). These territories are themselves subdivided into sub-nuclei distinguished by gene signatures and innervation pattern (Wagner, French and Veh, 2014; Wagner *et al.*, 2016; Juárez-Leal *et al.*, 2022), which have been largely corroborated by scRNA-seq analyses (Hashikawa *et al.*, 2020; Wallace *et al.*, 2020). In the zebrafish, two broad subdivisions, respectively dorsal (dHb) and ventral (vHb) of the habenulae have also been recognized and by their molecular signatures and projections respectively to interpeduncular nuclei and dorsal raphe, they are considered as homologous to the MHb and LHb components identified in mammals (Amo *et al.*, 2010; Beretta *et al.*, 2012; Okamoto and Aizawa, 2013). ScRNA-seq analysis yield support to this relationship and additionally highlighted the complexity of habenular neuronal populations, with the detection of 15 cell clusters, including 4 ventral and 10 dorsal ones (Pandey *et al.*, 2018). However, the number of signature markers supporting the correspondence between the dorsal and medial, or and ventral and lateral habenulae of teleosts and mammals remains low (Hashikawa *et al.*, 2020). Furthermore, the presence of similarities between the neuronal subpopulations within each territory remains elusive. Along the same line, a remarkable specificity of the habenulae is that they exhibit asymmetries, albeit highly variable in their degree, in many vertebrate species (Concha and Wilson, 2001). This trait exemplifies the difficulties encountered when comparing the mouse and zebrafish. The zebrafish habenulae exhibit asymmetries, which have been extensively characterized (Gamse *et al.*, 2003; Kuan *et al.*, 2007; Bianco and Wilson, 2009; deCarvalho *et al.*, 2014; Wang *et al.*, 2021) and have no equivalent in mammals, even though subtle size and shape differences between the right and the left habenulae have been detected in the latter (Ahumada-Galleguillos *et al.*, 2017; Hitti *et al.*, 2022). These difficulties may be related to lineage specific diversifications, obscuring similarities reflecting conserved ancestral features of organ organization. A classical strategy to test this hypothesis relies on the addition of an outgroup to comparisons, in order to bring to light similarities possibly lost in either lineage but retained in the other one, and infer ancestral versus divergent, lineage-specific traits. This strategy, using a chondrichthyan, the catshark *Scyliorhinus canicula*, as external reference, was successful to reveal ancestral asymmetries in gnathostomes and suggest an evolutionary scenario for diversifications (Lanoizelet et al. ms en prep.: Cf chapitre 1.). In order to expand this analysis to the general organization of habenulae, we have generated a genome-wide 3D RNA profile of the catshark habenulae. These data provide a refined view of their subdomain organization, which reveals dual neuronal identities reflecting both asymmetric and symmetric, time-dependent regulations and should be a reference for cross-species comparisons.

## 2. MATERIALS AND METHODS

### 2.1. Explant dissection, sectioning and RNA extraction from sections

Eggs from the catshark S. canicula were obtained from the Aquariology Service of the Banyuls sur Mer Oceanological Observatory. Juveniles (Ballard, Jean and Lechenault, 1993) were manually dissected and the habenulae explant was transferred to O.C.T. medium (Tissue-Tek O.C.T. compound, Sakura Finetek), frozen in liquid nitrogen prior to sectioning (16 μm cryostat sections). Each frozen section was transferred into Trizol (200 μL) immediately after collection. Total RNA was extracted using standard Trizol extraction protocol and purified on Macherey Nagel NucleoSpin RNA XS columns according to the manufacturer’s instructions.

### 2.2. llumina library construction and sequencing

cDNA synthesis, amplification, and Illumina library construction were conducted following (Holler and Junker, 2019) with the following modifications. Superscript II reverse-transcriptase was replaced by SuperScript IV (Invitrogen) and cDNA clean-up was done using Omega Bio-Tek’s Mag-Bind TotalPure NGS. Sequencing was conducted on the DNB-seq platform (100 bp paired-end).

### 2.3. Read mapping and counting

Reads were mapped onto the reference database of predicted genes. Briefly, the construction of this database (to be reported elsewhere) involved building an isoform collapsed version of the NCBI gene predictions annotated from the catshark genome assembly1 and a subsequent extension of 3′ UTRs. The resulting gene models are referred to as genes hereafter for simplicity. Reference indexing, read mappings and quantifications were done using Kallisto v0.44.0 using the –bias parameter and bootstrapping 100 times. Read counts were aggregated for each gene and each sectioning plane. Read counts per section were then treated by Spline Smoothing algorithm (Supplementary Figure 1).

### 2.4. 3D model construction

To generate a 3D binary mask, a juvenile habenulae explant dissected as described above was fixed in PFA 4%, DAPI stained, dehydrated in methanol and cleared in Benzyl Alcohol/Benzyl Benzoate (1/2) prior to mounting and imaging with a SP8X confocal microscope (Leica). A 3D binary mask was built using Fiji to obtain a binary image of the tissue. It was then reoriented transversally using a custom MATLAB script. The digital mask was locally modified to fit with 1D profiles of known genes and resized to match sectioning planes and section numbers along all three planes. The 3D expression genome-wide profile was reconstructed from spline smoothed read counts for each gene and each section plane using the MATLAB code reported in (Junker *et al.*, 2014). In short, following the virtual partitioning of the binary mask in serial digital sections as described above, an iterative process was applied to 1D profiles of each gene in each sectioning plane, multiplying the 3D expression of the gene by said profile in each plane in succession. 3D profiles of selected genes were then visualized using Fiji.

### 2.5. Statistical analysis

For the autocorrelation analysis, Moran’s indexes were calculated for each gene on a volume defined as the voxels on adjacent sections on all three section planes, giving a 3×3×3 cube centered on each voxel examined. Supports for the indexes thus obtained were estimated by statistical tests, consisting in calculating the Pearson correlation between the expression of a gene between all possible pairs of neighboring voxels, using the R libraries spdep (especially moran.test) and ade4. Pearson correlations between the 3D expressions of each possible pair of genes in each voxel (16 μm cube at the intersection of one section of each sectioning plane) were calculated using the R command cor with default parameters.

### 2.6. In situ hybridization of sections

ISH of paraffin sections were conducted using standard protocols as described previously (Lagadec *et al.*, 2015).

## 3. RESULTS

### 3.1. Generating genome-wide gene expression traces along AP, DV and LR axes

In order to obtain a digital transcriptomic map of the catshark habenulae, we manually dissected the organ from three juveniles and cryo-sectioned these explants along L/R (left/right), D/V (dorso-ventral) and A/P (antero-posterior) axes (Supplementary Figure 2A). This yielded a total of 87 sagittal, 45 horizontal and 48 transverse 16μm serial frozen sections (Supplementary Figure 2B). Each individual section was submitted to RNA extraction and cDNA synthesis using barcoded primers, which allowed to pool cDNA samples for linear amplification, Illumina library construction and sequencing (Supplementary Figure 2C). We thus obtained a total of 147.8, 52.3, and 86.2 million reads (PE-100) for sections along L/R, D/V and A/P axes respectively ( Supplementary Figure 3A,B,C). This corresponds to approximately 1.59M reads per section but a substantial heterogeneity was observed, in part due to differences in the amount of tissue in the section. For the bioinformatic analysis, reads were demultiplexed and mapped onto a gene reference obtained taking advantage of the catshark genome NCBI annotation and of transcriptomic resources available in our laboratory (Mayeur *et al.*, 2021). This allowed the generation of transcriptomic expression profiles for any predicted gene along each section plane. Those obtained for genes previously identified by a transcriptomic search for asymmetries (Lanoizelet et al. ms en prep.: Cf chapitre 1) are consistent with their broad expression characteristics, as inferred by ISH (Figure 1). Along the LR axis, read counts for *ScSox1* reach the highest levels in sections located to the left of the midline but are also detected above basal levels in a few right-sided successive sections (Figure 1A). Those for *ScKiss1* peak at the level of lateral-most right sections (Figure 1B), in line with the marked right enrichment observed in ISH. Similarly, along the AP axis, read counts for *ScStk32c* are high in anterior regions but return to basal levels posteriorly (Figure 1C), as expected from ISH profiles.

**Figure 1.**
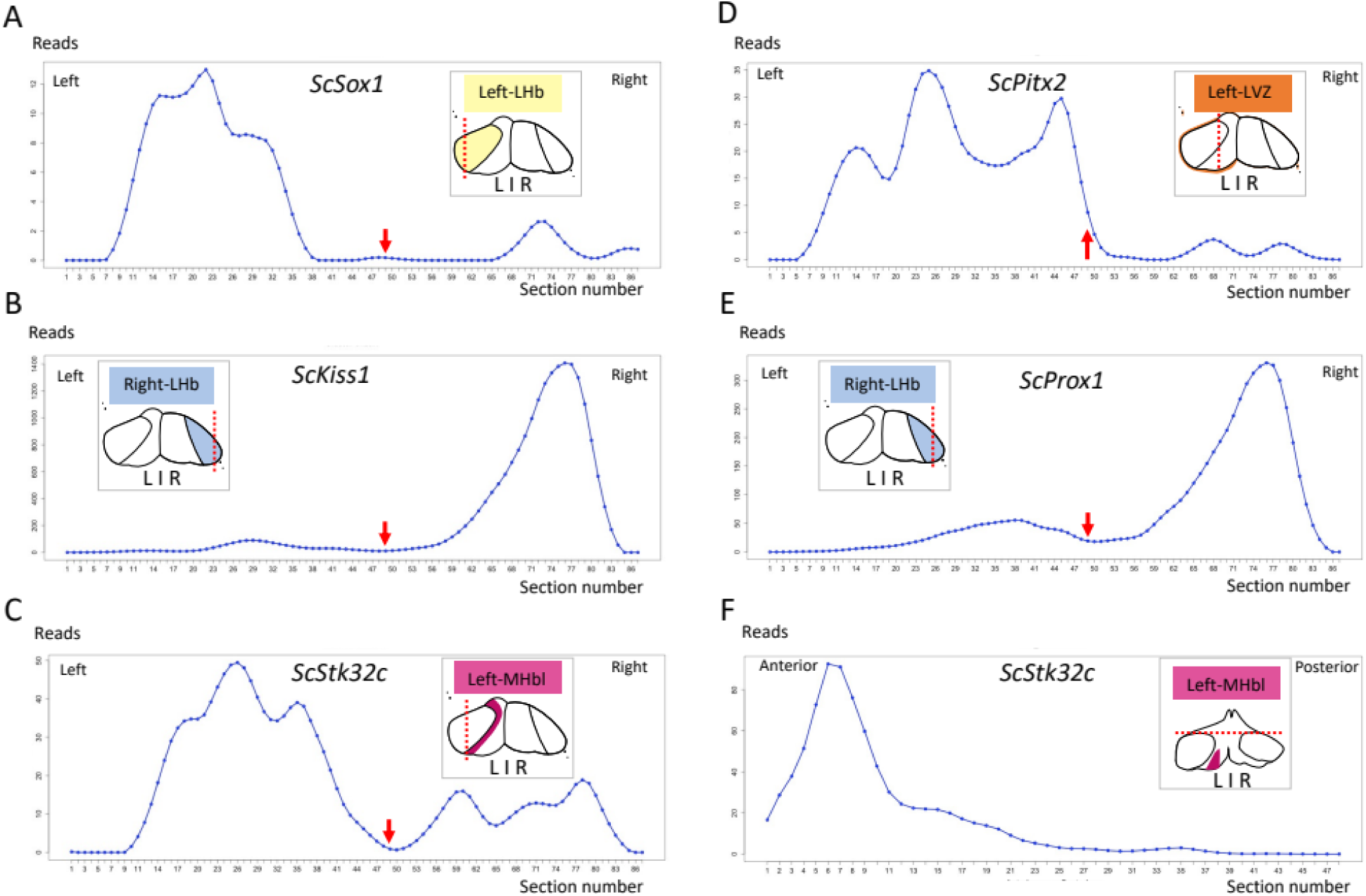
Expression traces for selected genes along sagittal (A-E) and transverse (F) planes. For each graph, section numbers and corresponding read counts are indicated along x- and y-axes, respectively. Sections 1-87 and 1-48 are respectively numbered from left to right in (A-E) and anterior to posterior in (F). Gene names are indicated at the top of each profile. For each profile, schemes in gray boxes represent the gene expression on front (A-E) or dorsal views of habenulae (F). Dotted red lines indicate section planes. The red arrow in (A-E) points to the level of midline of habenulae. Abbreviations: L/R, Left/Right.

**Figure 2.**
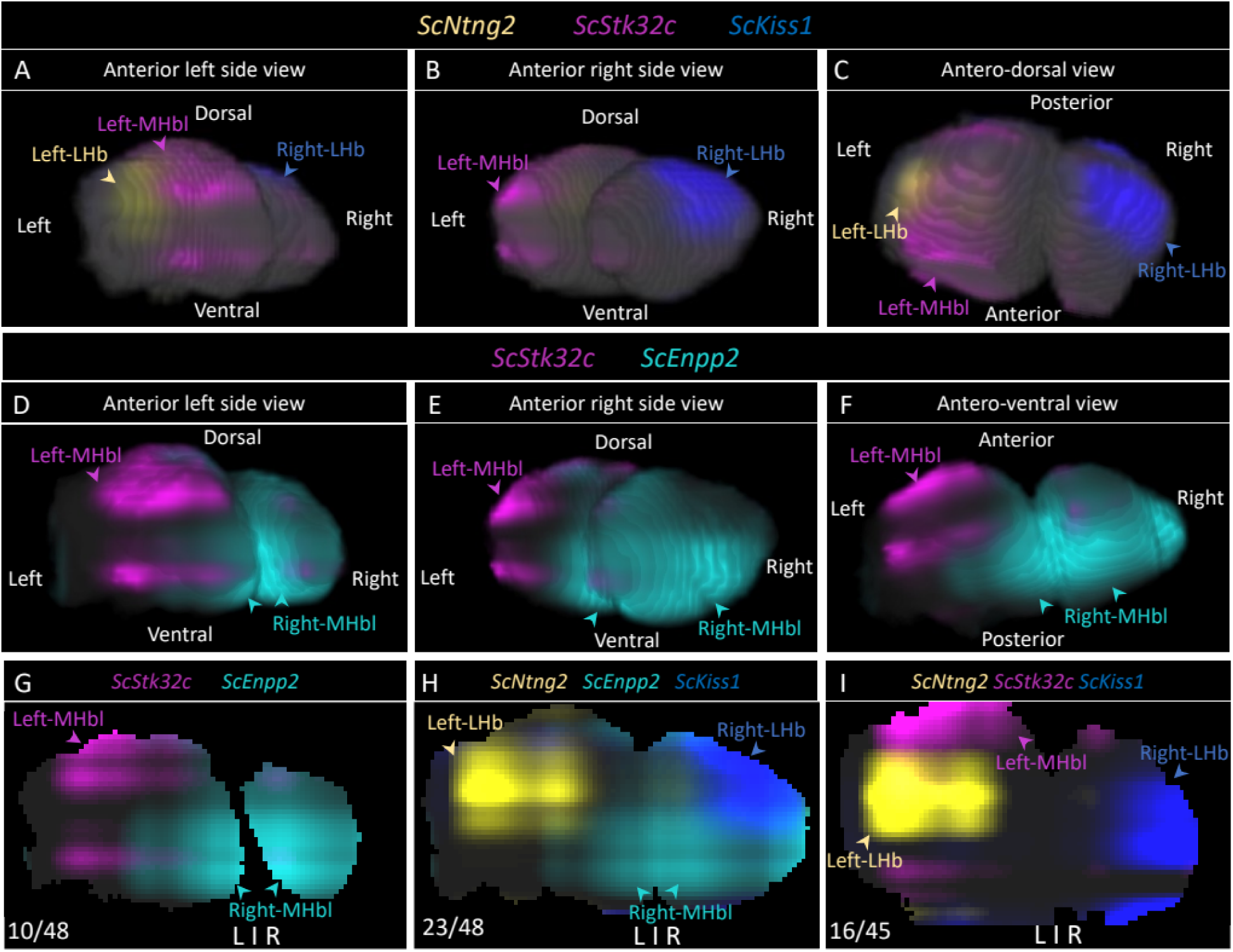
Digital 3D profiles of candidate genes. **(A-C)** show the combination of digital 3D profiles for *ScNtng2* (yellow)*, ScStk32c* (magenta) and *ScKiss1* (blue), with (A), (B) and (C) from anterior left side, anterior right side and antero-dorsal views respectively. **(D-F)** show the combination of digital 3D profiles for *ScStk32c* (magenta) and *ScEnpp2* (light blue) from anterior left side (D), anterior right side (E) and antero-ventral (F) views. **(G-H)** show digital transverse sections with digital profiles for *ScStk32c* (magenta) and *ScEnpp2* (light blue) in (G), *ScNtng2* (yellow)*, ScEnpp2* (light blue) and *ScKiss1* (blue) in (H). **(I)** shows digital horizontal section with digital profile for *ScNtng2* (yellow), *ScStk32c* (magenta) and *ScKiss1* (blue). Numbers in the left bottom corner in (G-I) refer to the numbering of the digital section shown within the series, from anterior to posterior (G-H) and dorsal to ventral for (I). Colored arrowheads point to digital territories corresponding to Left-LHb (yellow), Left-MHb (magenta), Right-MHb (light blue), Right-LHb (blue).

**Figure 3.**
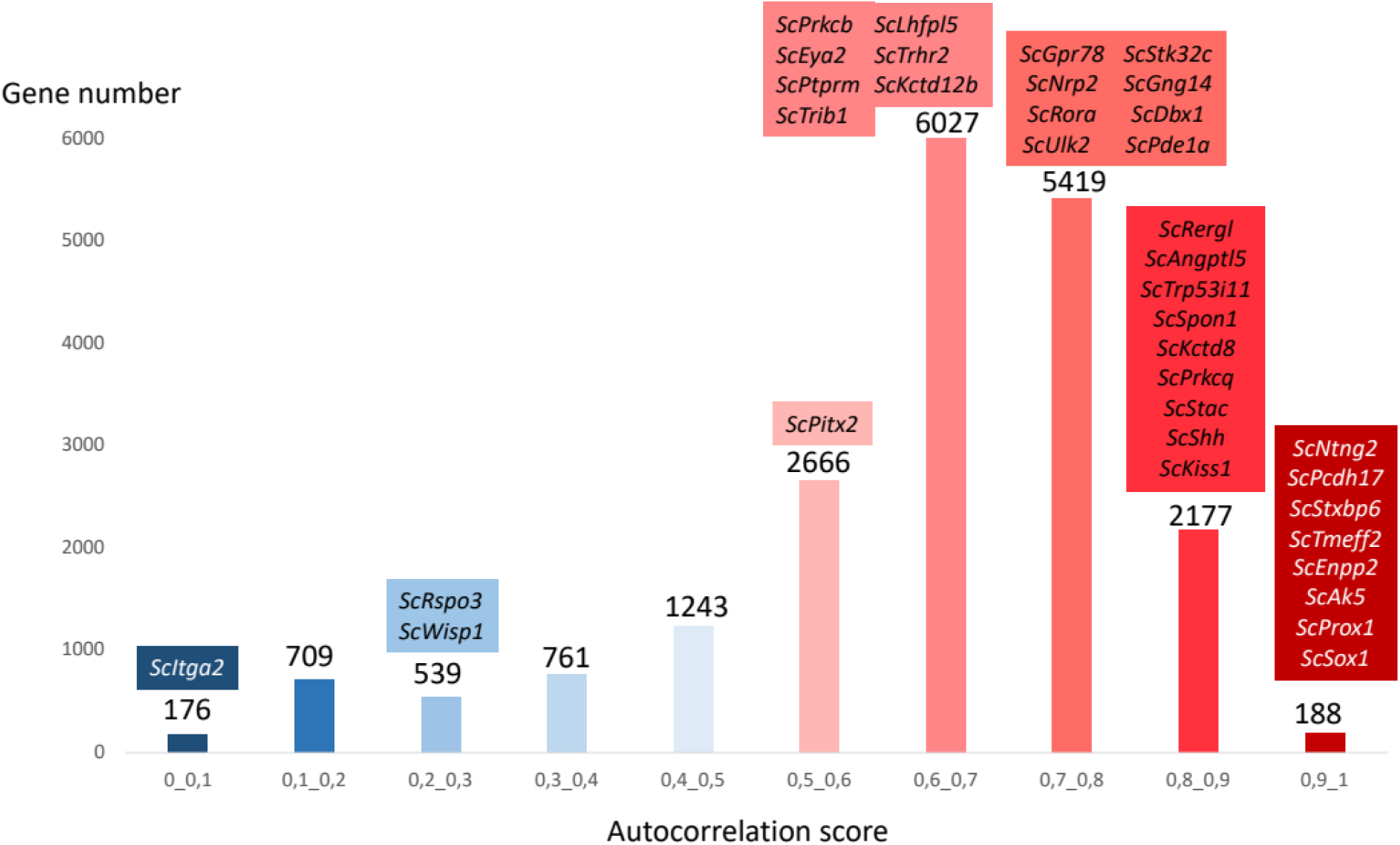
Distribution of genes depending on autocorrelation score. The bar plot shows the number of genes (y-axis) harboring autocorrelation scores within the range shown in x-axis. Bars are colored from the lowest autocorrelation scores (dark blue) to the highest ones (dark red). Genes previously characterized by ISH and known to harbor regionalized expressions in the habenulae are indicated and placed on the plot according to their autocorrelation score.

### 3.2. Generation of a 3D RNA model reflecting expression restrictions of habenula subdomain markers

In order to integrate the RNA profiles obtained along each axis into a 3D model, we projected the sequence data obtained for each section onto a digital mask generated using confocal microscopy (Supplementary Figure 2E,F), as described in (Junker *et al.*, 2014). This resulted in a 3D RNA profile, containing digital expression information for each individual gene model of the reference database (i.e. 30 270 gene models) and in each one of the 95892 voxels, identified by its (x, y, z) coordinates in the mask. In order to test this model and assess whether virtual expression patterns reflect expected ISH expression profiles, we focused on signature markers of the main habenula subdomains previously identified (Lanoizelet et al. ms en prep.: Cf chapitre 1). All selected genes exhibit regionalized expressions along the AP (*ScStk32c*) and LR (left-right) axis (*ScNtng2, ScKiss1, ScEnpp2*) and the corresponding digital territories are expected to occupy defined sectors, identified by distinctive (x,y,z) coordinates of the 3D habenula model. The broad expression restrictions of these genes along the axes, i.e. anterior restriction and left enrichment of *ScStk32c*, right enrichment of *ScEnpp2*, left and right restrictions of *ScNtng2* and *ScKiss1* respectively, are retrieved in the 3D model (Figure 2). While digital profiles do not reproduce accurate territory shapes or expression details such as the strict complementarity between lateral and medial territories, those obtained for each one these four genes harbor distinctive features, not shared by the other ones, thus defining digital versions of previously identified habenula territories (Left- and Right-LHb; Left-MHbl; Right-MHbl).

### 3.3. Identification of habenular territories by autocorrelation analysis

In order to identify novel habenular territories, we conducted a systematic search for genes of regionalized digital profile, using autocorrelation as a proxy for digital expression regionalization (Mayeur *et al.*, 2021). Autocorrelations scores (A) were calculated for all gene models, which were ranked accordingly (Supplementary Table 2). Most of the genes previously analyzed and known to exhibit regionalized expressions (Lanoizelet et al. ms en prep.: Cf chapitre 1) show high autocorrelation scores (A>0.6 for 32/37 genes, Figure 3). Among the four exceptions (*ScPitx2*, *ScItga2*, *ScRspo3*, *ScWisp1*), the three former ones are expressed in a thin neural progenitor layer, referred to lateral ventricular zone (LVZ, Lagadec et al., 2018), and they are largely restricted to the left, in line with the profile of read counts observed along the LR axis (Supplementary Figure 4). Their 3D profiles are unlikely to yield high autocorrelation scores by their geometry (Supplementary Figure 4). Concerning the latter (*ScWisp1*), very low read counts were obtained, which leads to a decrease in autocorrelation values and support (Mayeur *et al.*, 2021). Conversely, a high proportion of the gene models show high autocorrelation scores (7784 genes with A>0.7; 11446 with A>0.6), suggesting a rich repertoire of marker genes for discrete habenular subdomains (Figure 3). To gain insight into these territories, we focused on the 50 gene models exhibiting the highest autocorrelation scores (A>0.67), restricting the analysis to annotated ones (i.e. 42 genes). Among those, four were already identified as subdomains markers (*ScNtng2*, A=0.99; *ScPcdh17*, A=0.96; *ScStxbp6*, A=0.95; *ScEnpp2*, A=0.93). A systematic ISH characterization on juvenile habenula sections was conducted on the 40 remaining ones (Figure 4). This analysis first highlighted novel signature markers of previously known territories (Figure 4A), including Left-LHb (Figure 4A, Suppl. Table 2) (*ScTmem65*, A=0.99; *ScCntnap1*, A=0.95; *ScStmn2*, A=0.93), Right-LHb (Figure 4A, Suppl. Table 2) (*ScSez6l,* A=0.95; *ScSt6galnac5,* A=0.94; *ScPldxdc1*, A=0.93), MHb (Figure 4A, Suppl. Table 2) (*ScGng8*, A=0.95; *ScCbarp,* A=0.93), Left-MHbl (Suppl. Table 2) (*ScFabpl,* A=0.93) and Right-MHbl (Figure 4A, Suppl. Table 2) (*ScCplx3,* A=0.95; *ScCamkV,* A=0.93). More unexpectedly, we also identified genes sharing expressions in the left and right lateral habenulae ( *ScGap43*, A=0.94; *ScVwab5b1,* A=0.93, Figure 4B1,B2), as well as in left and right components of the lateral part of the medial habenula (*ScPitpnm3*, A=0.93, Figure 4B3) and in the complementary medial part of the medial habenula (*ScPhlda2,* A=0.96, Figure 4B4). The corresponding territories were therefore referred to as LHb, MHbl and MHbm hereafter. Finally, we found evidence for a partitioning of Left- and Right-LHb, with restricted expressions of *ScP65b* and *ScNefl,*(A=0.93/0.92; dorsal subdomain of Left-LHb, Figure 4C2-C3), *ScCsrp1, (*A=0.93; ventral subdomain of Right-MHbl, Figure 4C1), *ScAlpl* (A=0.93; dorsal subdomain of Right-LHb, Figure 4C4). Finally different combinations of the territories identified above were observed (*ScTmeff2*, A=0.95; *ScBcat1,* A=0.94; *ScNpb,* A=0.94; *ScPenk,* A=0.94; *ScDcaf7,* A=0.94; *ScNptxr,* A=0.93, Figure 4D1, D2, D3 and D4).

**Figure 4.**
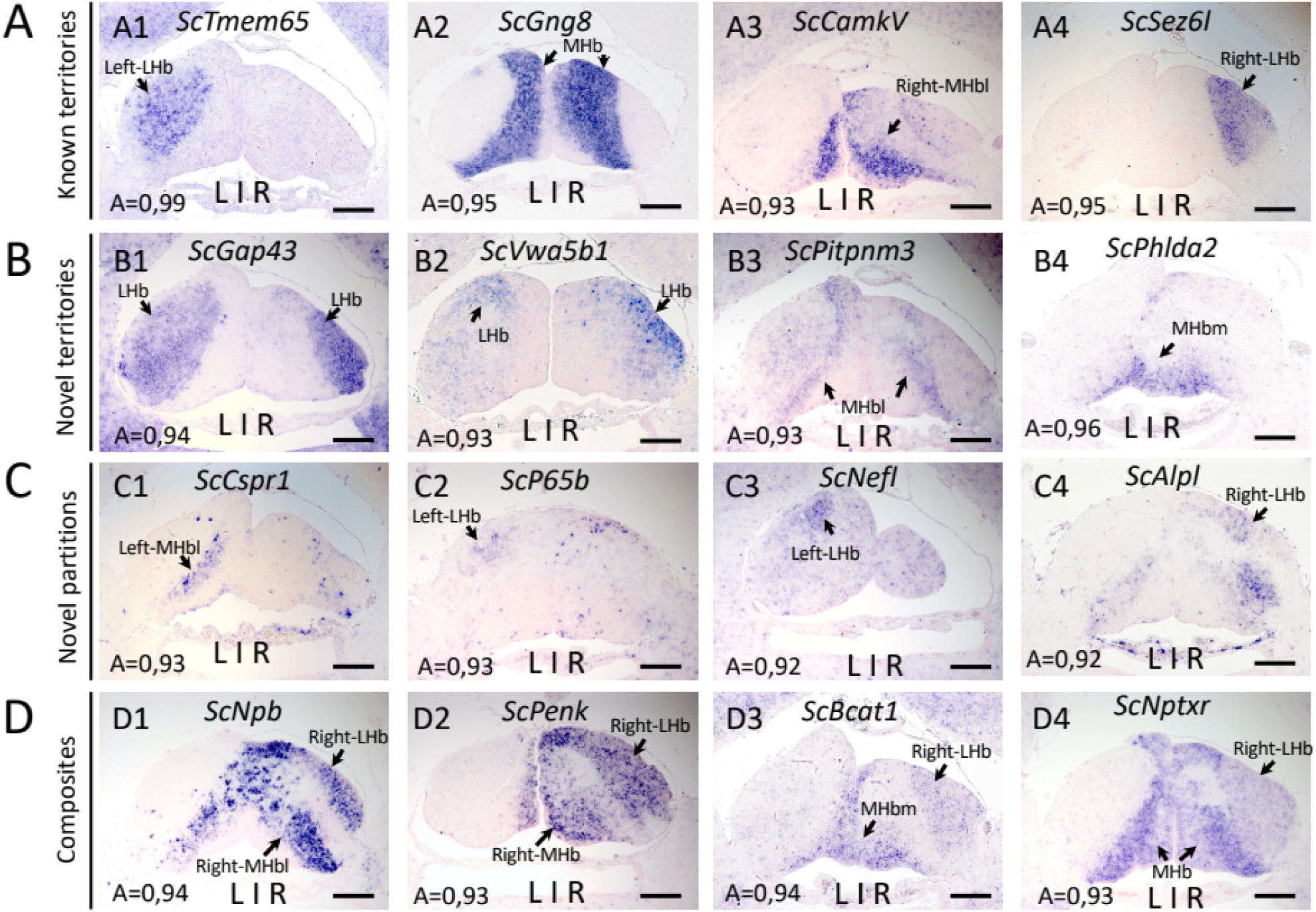
Expression profiles of selected genes harboring high autocorrelation scores. **(A-D)** show expression profiles corresponding respectively to previously characterized territories (A), novel ones (B), subdomains of the former two (C), combinations of the former two (D). **(A1-A4,B1-B4,C1-C4,D1-D4)** show transverse sections of juvenile catshark habenulae after ISH with probes for *ScTmem65* (A1), *ScGng8* (A2), *ScCamkV* (A3), *ScSez6l* (A4), *ScGap43* (B1), *ScVwa5b1* (B2), *ScPitpnm3* (B3), *ScPhlda2* (B4), *ScCspr1* (C1), *ScP65b* (C2), *ScNefl* (C3), *ScAlpl* (C4), *ScNpb* (D1), *ScPenk* (D2), *ScBcat1* (D3), *ScNptxr* (D4). For each gene the autocorrelation score is shown in the bottom left corner. Labeled territories are identified by thin arrows. Abbreviations: L/R, Left/Right; Scale bar = 200 μm.

### 3.4. Analyses of correlations with territory markers

In order to expand the marker repertoire of habenular territories, we conducted correlation analyses as described in (Mayeur *et al.*, 2021), focusing on the six major habenular territories identified above: Left-LHb, LHb, Right-LHb, Left-MHbl, MHb, Right-MHbl. For each one of these subdomains, we selected with two reference markers, exhibiting strongly autocorrelated digital profiles, validated by ISH analysis (respectively *ScNtng2*/*ScSox1*, *ScGap43*/*ScVwab5b1*, *ScStxbp6*/*ScKiss1*, *ScSpon1*/*ScStk32c*, *ScGng8*/*ScKctd8*, *ScEnpp2*/*ScCamkv*), and calculated correlation scores with these markers for all gene models (Supplementary Table 3 and 4). For each territory, results obtained with the two reference markers selected yield coherent results, genes harboring high (respectively low) correlation scores with either one similarly harboring high (respectively low) correlation scores with the other one (Supplementary Figure 6). Conversely, correlation scores are inversely related between exclusive, left- or right-enriched domains, whether in LHb or MHb (Left-LHb versus Right-LHb, Supplementary Figure 7A; Left-MHbl versus Right-MHbl, Supplementary Figure 8A). As expected, this inverse relationship was less marked between overlapping territories (Left- or right-LHb versus LHb, Supplementary Figure 7B,C; Left- or right-MHb versus MHb, Supplementary Figure 8B,C). The same conclusion applies to comparisons of correlation scores between exclusive medial and lateral territories (Supplementary Figure 9). We also analyzed the distribution of gene models depending on correlation values with reference markers for each of the territories analyzed (respectively *ScNtng2*, *ScVwab5b1*, *ScKiss1*, *ScStk32c*, *ScGng8*, *ScEnpp2* for Left-LHb, LHb, Right-LHb, Left-MHbl, MHb, Right-MHbl) and indicated known markers, including those identified by the autocorrelation analysis described above, on these diagrams (Figure 5). In all cases, the peak of the distribution is found for correlation values comprised between -0.1 and 0.1, suggesting that signature markers for these territories only concern a small proportion of the genes. In all cases with one exception (correlation with Right-MHb, for which a single marker other than *ScEnpp2* was known), genes specifically expressed in one of the territories analyzed harbor correlation values>0.5 with the corresponding reference marker. Along the same line, genes showing correlation values >0.6 with a given territory marker never include known markers for other territories. Taken together, these data validate the correlation analysis as a means to identify novel territory markers, also suggesting that lists of genes with correlation values >0.5 should be significantly enriched in such markers.

**Figure 5.**
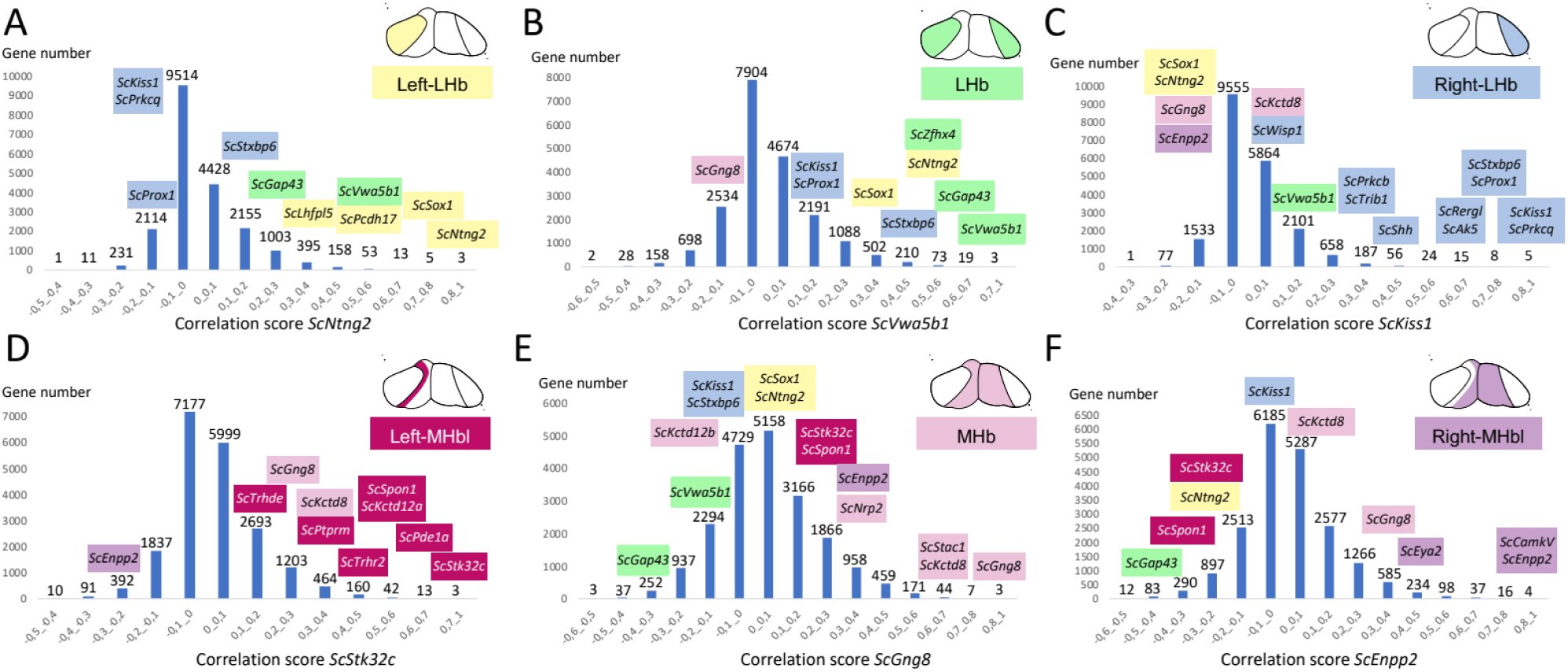
Distribution of genes depending on correlation scores. **(A-F)** show the number of genes (y-axis) harboring correlation scores within the range shown in x-axis with *ScNtng2* (A), *ScVwa5b1* (B), *ScKiss1* (C), *ScStk32c* (D), *ScGng8* (E) and *ScEnpp2* (F). Schemes in the upper left quadrant of each bar plot show front views of the habenulae with the territory labeled by each of these reference markers in color (Left-LHb, yellow; LHb, green; Right-LHb, blue; Left-MHb, dark magenta; MHb, pink; Right-MHb, purple). Previously characterized markers of these territories are shown shaded with the same color code and are placed on the plot according to their correlation score.

**Figure 6.**
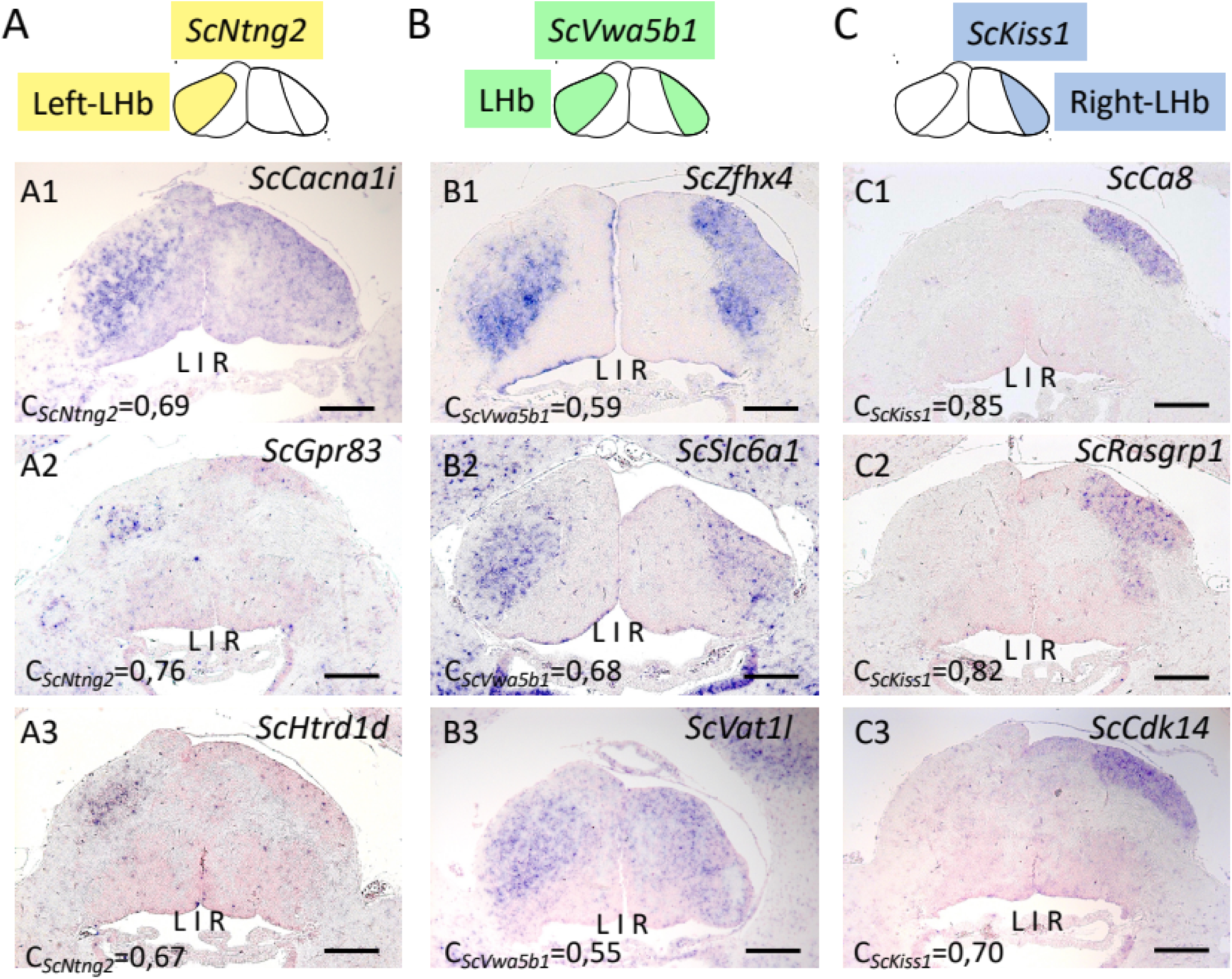
ISH expression profiles of novel markers of lateral habenula territories. **(A-C)** respectively show transverse sections of juvenile catshark habenulae for genes selected based on correlation scores with respectively *ScNtng2* (A), *ScVwa5b1* (B) and *ScKiss1* (C). The territories labeled by each of these three markers are schematized with Left-LHb in yellow, LHb in green and Right-LHb in blue. Expression is shown for *ScCacna1i* (A1), *ScGpr83* (A2), *ScHtr1d* (A3), *ScZfhx4* (B1), *ScSlc6a1* (B2), *ScVat1l* (B3), *ScCa8* (C1), *ScRasgrp1* (C2) and *ScCdk14* (C3). For each gene, the correlation score with the reference marker taken into account for its selection is shown in the bottom left corner. Scale bar = 200 μm.

**Figure 7.**
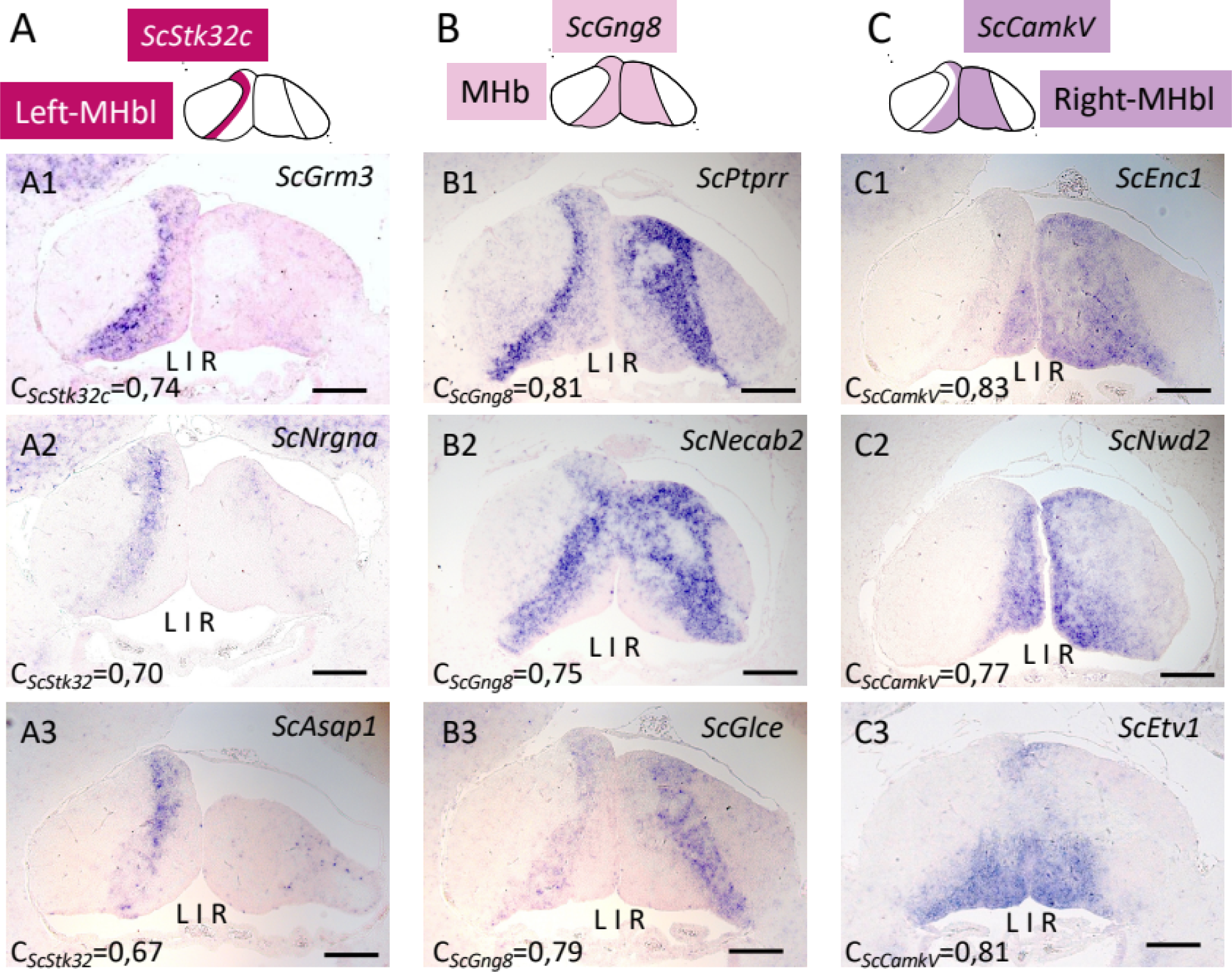
ISH expression profiles of novel markers of medial habenula territories. **(A-C)** respectively show transverse sections of juvenile catshark habenulae for genes selected based on correlation scores with respectively *ScStk32c* (A), *ScGng8* (B) and *ScCamkV* (C). The territories labeled by each of these three markers are schematized with Left-MHbl in dark red, MHb in light purple and Right-MHbl/MHbm in dark purple. Expression is shown for *ScGrm3* (A1), *ScNrgna* (A2), *ScAsap1* (A3), *ScPtprr* (B1), *ScNecab2* (B2), *ScGlce* (B3), *ScEnc1* (C1), *ScNwd2* (C2) and *ScEtv1* (C3). For each gene, the correlation score with the reference marker taken into account for its selection is shown in the bottom left corner. Scale bar = 200 μm.

**Figure 8.**
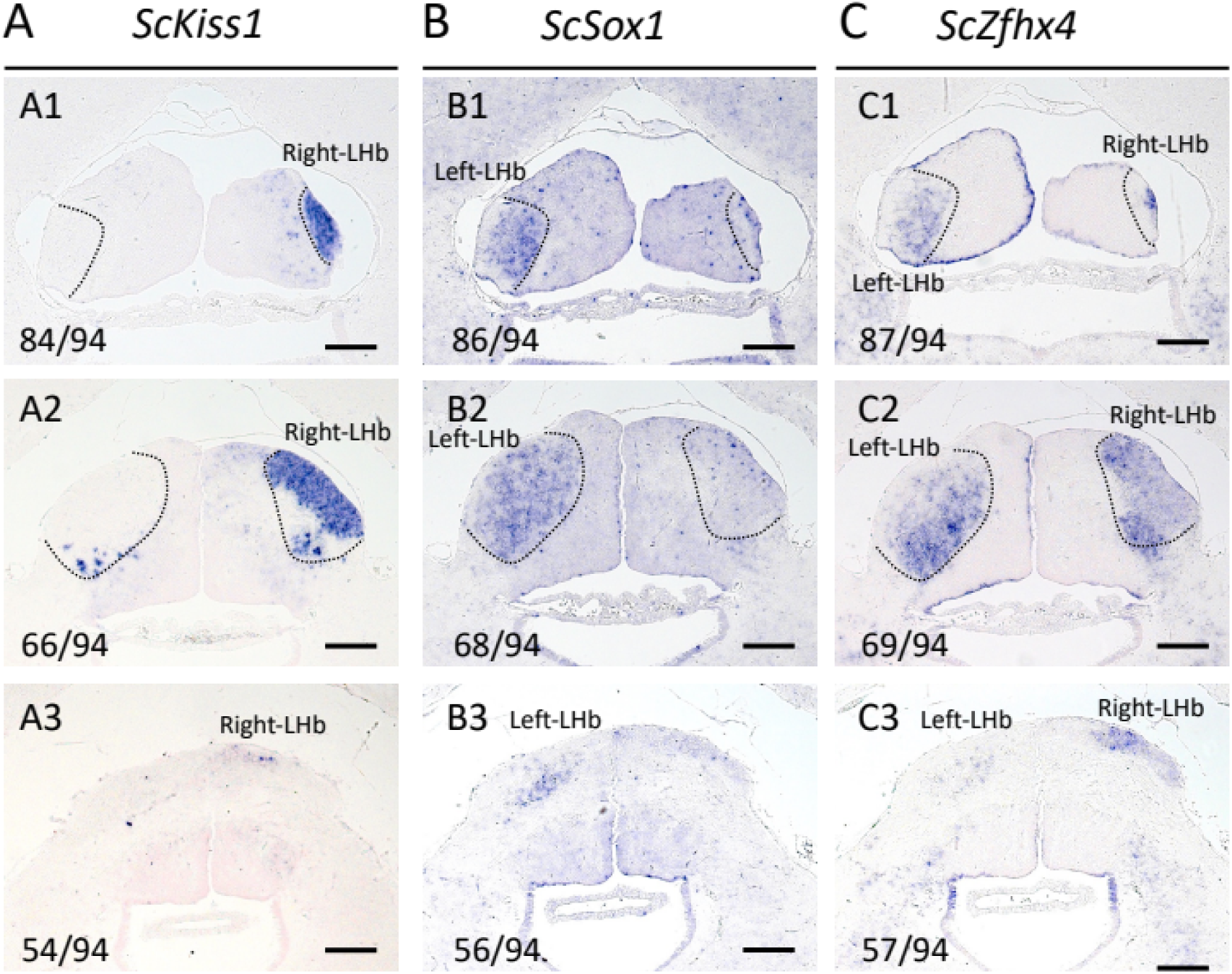
Comparison between ISH profiles of Left-LHb, Right-LHb and LHb markers. **(A-C)** show transverse sections of habenulae from the same individual (juvenile) after ISH with probes for *ScKiss1* (A), *ScSox1* (B) and *ScZfhx4* (C). Sections are shown at 3 different levels: anterior (A1,B1,C1), medial (A2,B2,C2) and posterior (A3,B3,C3). The numbering of sections is shown in the bottom left corner from the posterior-most section (1) to the anterior-most one (94). Scale bar = 200 μm.

**Figure 9.**
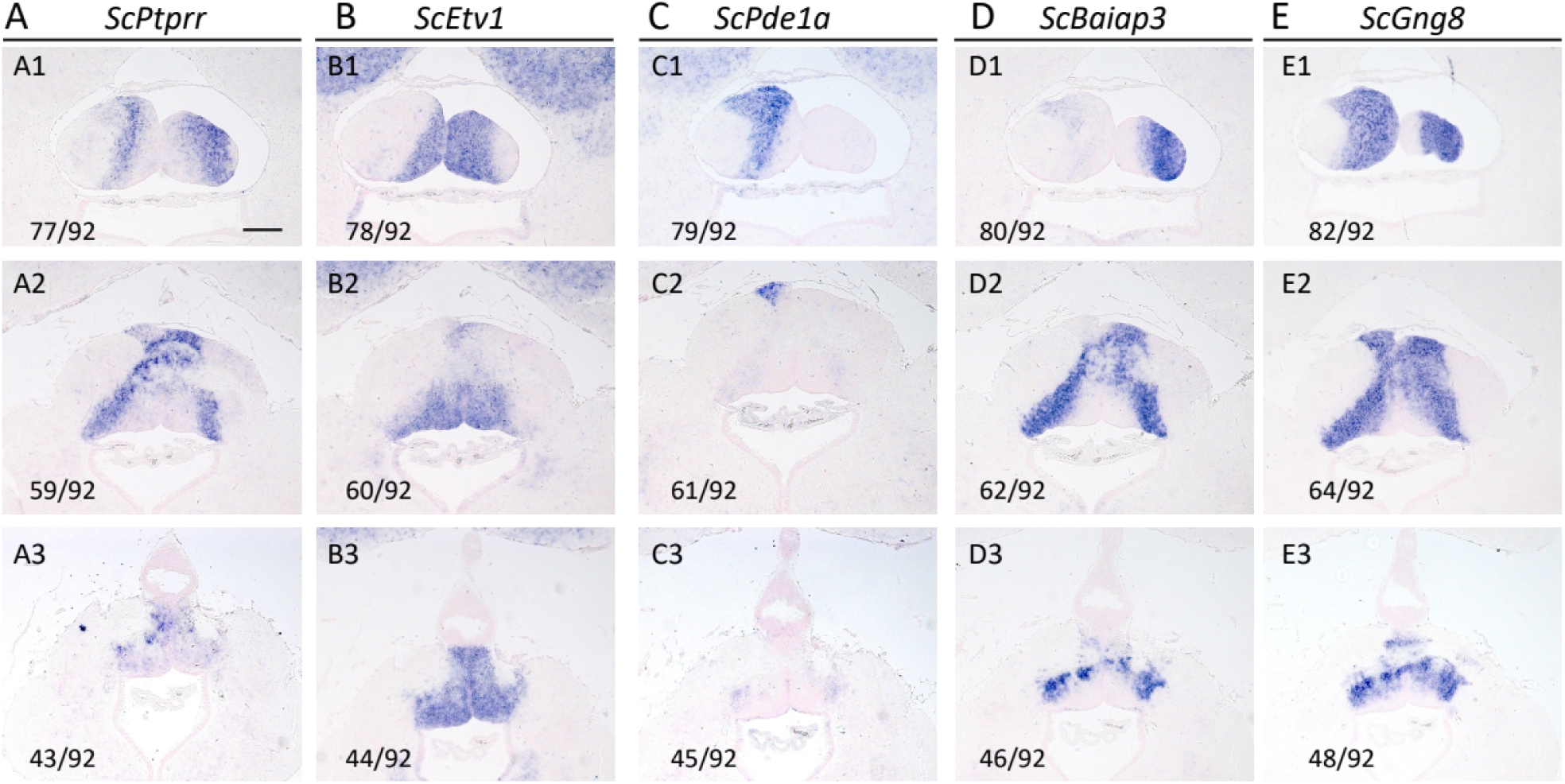
Comparison between ISH profiles of Left-MHbl, Right-MHbl, MHbl and MHbm markers. **(A-E)** show transverse sections of habenulae from the same individual (juvenile) after ISH with probes for *ScPtprr* (A), *ScEtv1* (B), *ScPde1a* (C), *Sc Baiap3* (D) and *ScGng8* (E). Sections are shown at 3 different levels: anterior (A1,B1,C1,D1,E1), medial (A2,B2,C2,D2,E2) and posterior (A3,B3,C3,D3,E3). The numbering of sections is shown in the bottom left corner from the posterior-most section (1) to the anterior-most one (92). Scale bar = 200 μm.

### 3.4. Expanding the marker repertoire for habenula broad territories

We next took advantage of the lists of genes correlated to subdomain markers (respectively *ScNtng2, ScVwa5b1, ScKiss1*, *ScStk32c*, *ScGng8* and *ScEnpp2/ScCamkv* for Left-LHb, LHb, Right-LHb, Left-MHbl, MHb and Right-MHbl) to select novel candidates (correlation scores > 0.5) for further ISH analysis. Concerning genes selected on the basis of correlation to markers of lateral territories ( Figure 6, Supplementary Table 3), ISH characterizations led to the identification of novel markers for Left-LHb (*ScCacna1i* : Figure 6A1), LHb (*ScSlc6a1, ScZfhx4, ScVat1l*: Figure 6B1-3) and Right-LHb (*ScCa8, ScRasgrp1, ScVwc2l, ScCdk14, ScLrrn3, ScFrrs1 and ScBtg1;* Figure 6C1-3, Supplementary Figure 5A).This analyses also resulted in the identification of genes expressed in a dorsal subterritory of Left-LHb (*ScGpr83, ScHtr1d*: Figure 6A2-3) and Right-LHb (*ScC2orf66* : Supplementary table 3, Supplementary Figure 5C). Concerning genes selected on the basis of correlation to markers of medial territories (Figure 7, Supplementary Table 4), ISH characterizations led to the identification of novel markers for Left-MHbl (*ScGrm3, ScNrgna, ScPmp22, ScAsap1, ScMap7d1, ScMpp2, ScSstr2*; Figure 7A1-3, Supplementary Table 4, Supplementary Figure 5), MHb (*ScNecab2*; Figure 7B) and Right-MHbl (*ScEnc1, ScIna, ScRtn4rl1, ScBaiap3* and *ScNwd2;* Figure 7C1-C3, Supplementary Figure 5B). This analyses also yielded genes exhibiting regionalized expressions in subdomains of the reference MHb territory, excluding either its medial or its lateral component, as observed for *ScPtprr*, *ScGlce* and *ScArhgdib* (novel MHbl markers; Figure 7B1,B3; Supplementary figure 5B), and *ScEtv1* (novel MHbm marker; Figure 7C3). Taken together, these data validate the possibility to identify novel markers of known territories, but also markers of subdomains included within these territories.

### 3.5. ISH characterization of the habenula medial to lateral organization

An unexpected result of the above analyses is the identification for the first time of genes exhibiting bilateral expressions in both left and right lateral habenula territories (Left- and Right-LHb), but also in both previously identified left and right lateral components of the medial habenula (Left- and Right-MHbl) as well as in the complementary medial component of MHb (MHbm). To confirm the relative organization of these left, right and bilateral components of the lateral and medial habenulae, we performed ISH on adjacent habenular sections of the same individual using markers for these lateral and medial territories (Figure 8 and Figure 9 respectively). This analysis confirms that the combination of Left-LHb and Right-LHb, detected by ISH with probes for *ScSox1* (Figure 8B) and *ScKiss1*(Figure 8A), is largely superimposable with LHb, expressing *ScZfhx4* (Compare Figure 8A, 8B with Figure 8C). Similarly, in the *ScGng8* positive medial habenula (MHb) (Figure 9E), the territory expressing *ScEtv1* (Figure 9B), a MHbm marker, is complementary to the one expressing *ScPtprr* (Figure 9A, compare Figure 9A1,A2 with 9B1,B2), a MHbl marker. The latter is largely superimposable to the combination of *ScPde1a* (Figure 9C) and *ScBaiap3* (Figure 9D) territories (Left- and Right-MHbl). This highlights a lateral to medial organization of habenulae, consisting of a succession of radial domains, which differ by their size but share common gene signatures between their left and right components.

## CONCLUSION AND PERSPECTIVES

A transcriptomic comparison between the catshark left and right developing habenulae had revealed a highly asymmetric subdomain organization, likely to reflect the gnathostome ancestral state (Mazan, pers. communication). Here, we have used RNA tomography to alleviate the bias of our previous characterization, restricted by its methodology to genes harboring asymmetric expressions. While this approach does not allow a high resolution due to different limitations -unavoidable differences between the four specimens required, imperfect control of sections planes, approximations of the iterative proportional fitting algorithm-, it is an efficient way to resolve regionalized territories along the section axis used (Mayeur *et al.*, 2021). The digital profile obtained provides a refined molecular map of this organ, including genome-wide, 3D expression data and allowing a discrimination between broad habenular territories. The use of correlation and autocorrelation analyses yields an expanded repertoire lists of their signature markers (Figure 10). These data therefore provide a novel reference for comparative analyses of habenula organization across gnathostomes.

**Figure 10.**
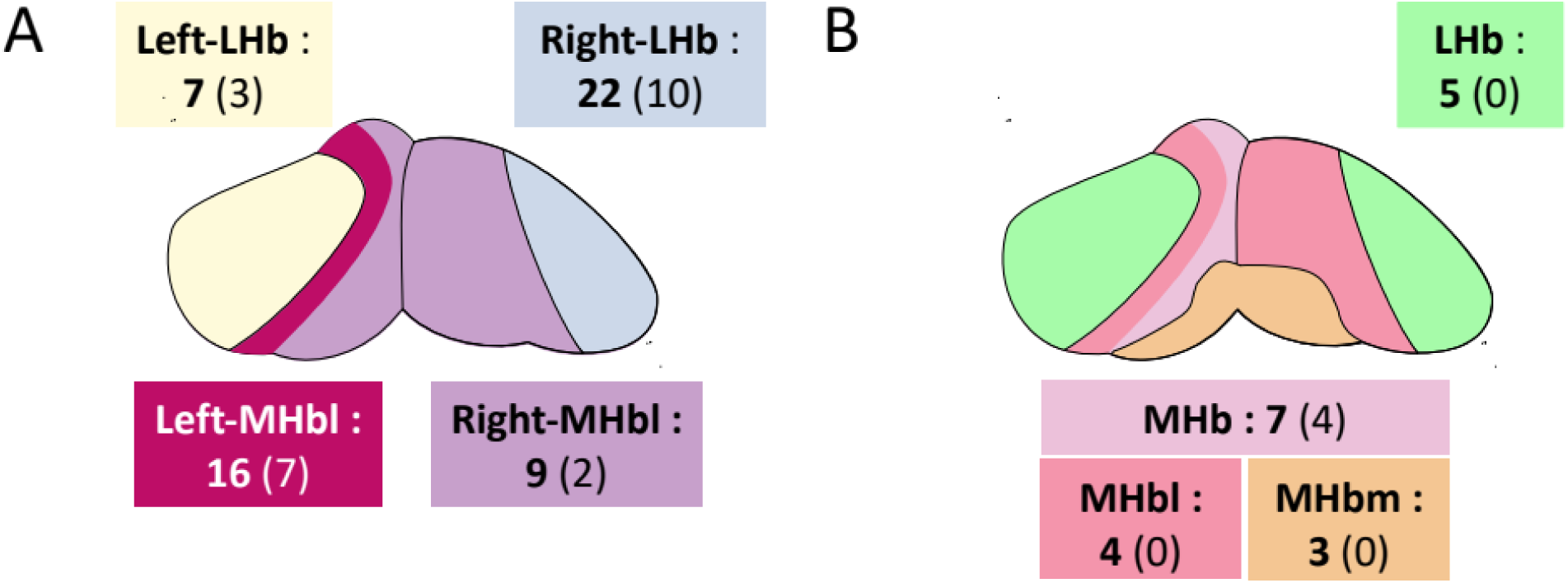
Novel catshark juvenile mediolateral organization of habenulae. (**A**) Asymmetric organization of habenulae. (**B**) Temporal-evoked mediolateral organization of habenulae. Bold numbers show the number of markers supporting each domain after the study while numbers in bracket states the number of markers before the study.

Several conclusions emerge from their analysis. A first one is that territories, harboring mirror locations on each side of the midline and thought to be highly asymmetric based on markers inferred from our initial characterization, share in fact highly specific molecular signatures. For instance, Left- and Right-LHb were characterized with extensively divergent molecular identities following our previous analysis, but the Tomo-seq characterization unveils an underlying unity, with several markers specifically expressed in these territories on both sides (*ScGap43*, *ScVwab5b1*, *ScZfhx4*). The same conclusion applies to left and right external domains of the medial habenula, which share expression of *ScPitpnm3, ScGlce, ScArhgdib* and *ScPtprr*. Together with the identification of an additional bilateral sub-territory complementary to the former in the medial part of MHb ( expressing *ScPhlda2, ScEtv1, ScNeurod1*), this reveals a lateral to medial organization in bilateral radial bands of defined molecular identity. This spatial pattern is fully consistent with a temporal regulation of neurogenesis and a construction of habenulae by successive additions of radial territories, from lateral-most to medial levels. This is reminiscent of the temporal regulation of neurogenesis and neuronal identity elaboration, as reported in the zebrafish habenula (Aizawa *et al.*, 2007; Fore *et al.*, 2020) or other brain regions (Nguyen *et al.*, 1999; Lein *et al.*, 2017). We suggest that the dual neuronal identities observed in these territories, with both bilateral and asymmetric features, reflects the combination of a symmetric temporal regulation, and an asymmetric one, which interacts with the former and differentially shape neuronal identities between the right and left habenula sides.

These data and hypothesis raise novel, mechanistic and evolutionary questions. From a mechanistic perspective, the mechanisms controlling the formation of the bilateral, medial to lateral pattern unveiled by this analysis remain completely unknown. The set of molecular markers identified provide handles to address the timing of its establishment and signaling pathways possibly involved using pharmacological approaches. They also provide reference data to explore the dynamic of intrinsic regulatory programs by analyzing cell trajectories from neural progenitors to differentiated neurons. Despite its limitations for gene loss-of-function approaches, the catshark should be an excellent model for such approaches by the spatial and temporal resolution that it allows during habenula development.

Concerning evolutionary aspects, systematic comparisons with the mouse and zebrafish remain to be conducted. While scRNA-seq and Tomo-seq data are difficult to integrate using bio-informatic tools, the catshark may provide a valuable reference for a candidate gene approach, due both to the outgroup position of chondrichthyans relative to sarcopterygians and actinopterygians and to the maintenance of ancestral asymmetry traits in this species (Michel, Lagadec and Mazan, ms in prep.).

## Supplementary Material

**Supplementary Figure 1.**
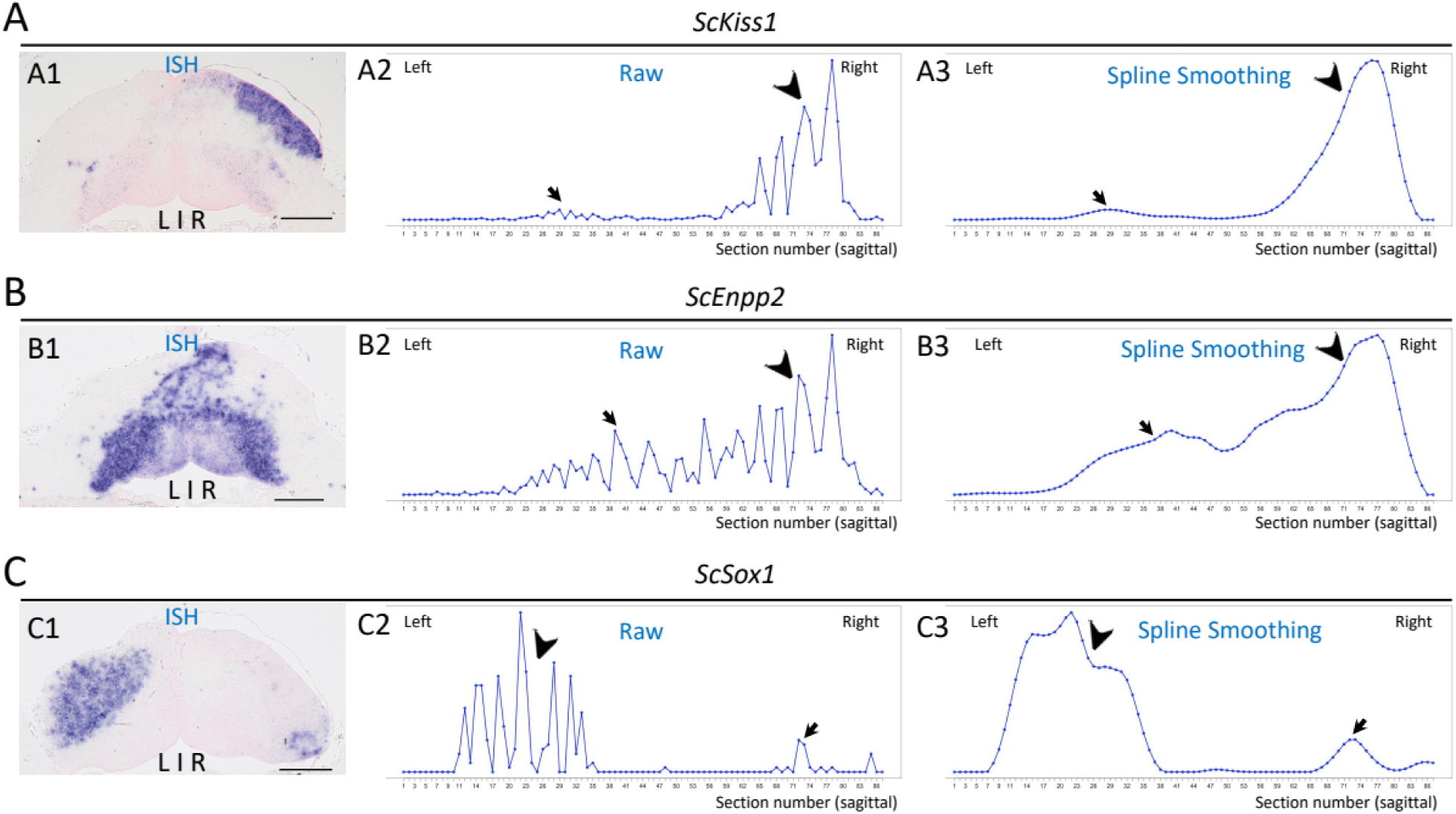
Effect of spline-smoothing on expression traces along one axis. **(A-C)** show the effect of spline-smoothing on expression traces of *ScKiss1*(A), *ScEnpp2* (B) and *ScSox1* (C) along the left-right axis. (A1,B1,C1) show transverse sections of catshark juvenile habenulae after ISH. (A2,B2,C2) and (A3,B3,C3) show read counts (y-axis) per section (x-axis) from left to right, respectively without (raw data) and with spline-smoothing treatment. Arrowheads and thin arrows respectively point towards major peaks over neighboring or adjacent sections and more minor signals, observed in ISH and reproduced in the digital data. The latter are maintained with the parameters used for spline-smoothing.

**Supplementary Figure 2:**
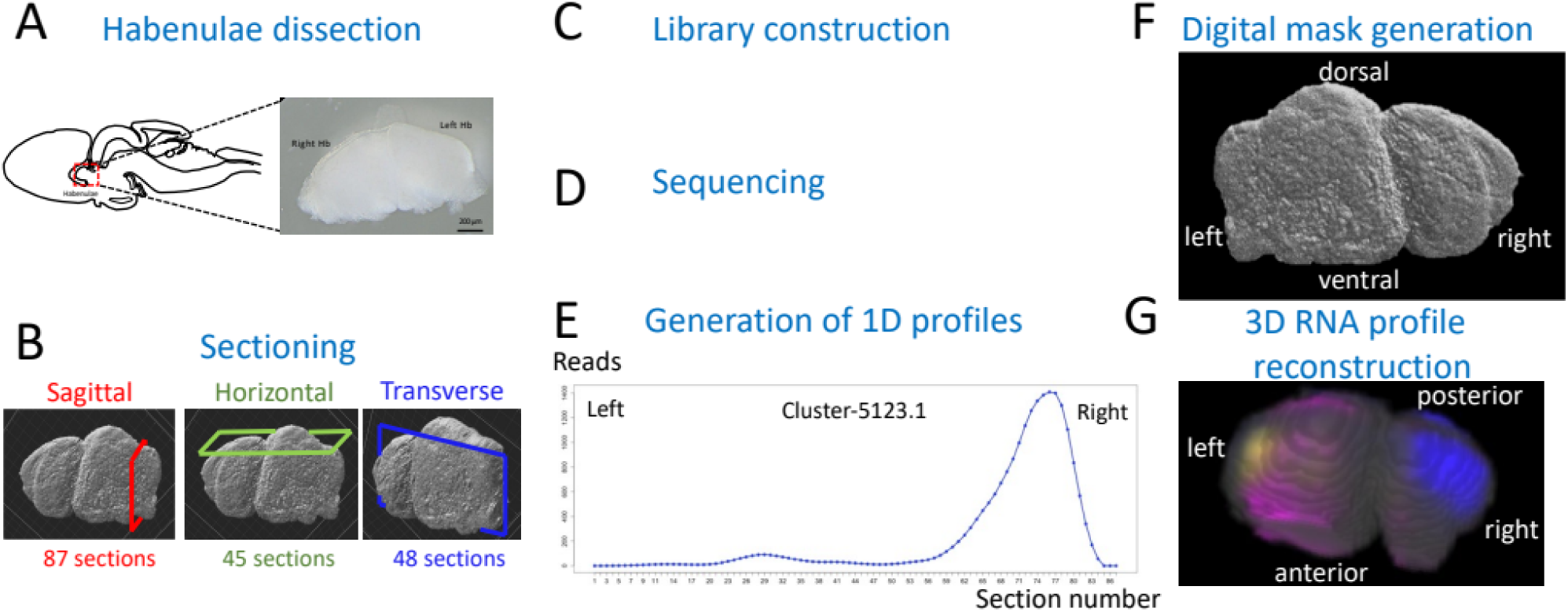
Main experimental steps used to generate a genome-wide 3D profile of the catshark habenulae. **(A)** Explant dissection: the scheme on the left shows a left side view of the catshark brain, with the habenula location boxed in red and the photograph on the left shows a posterior view of the dissected habenula explant. **(B)** Cryo-sectioning: schemes show the planes used to generate series of cryo-sections (sagittal, red; horizontal, green; transverse, blue). The total number of sections obtained along each plane is in E) Visualization of expression traces along each axis for all gene models. **(F)** Generation of a digitized model by confocal microscopy. **(G)** Generation of the genome-wide 3D RNA profile by computation of digital expressions for each voxel and gene model using the Integrative Proportional fitting algorithm. D, dorsal; V, ventral; Ant, anterior; Post, posterior.

**Supplementary Figure 3.**
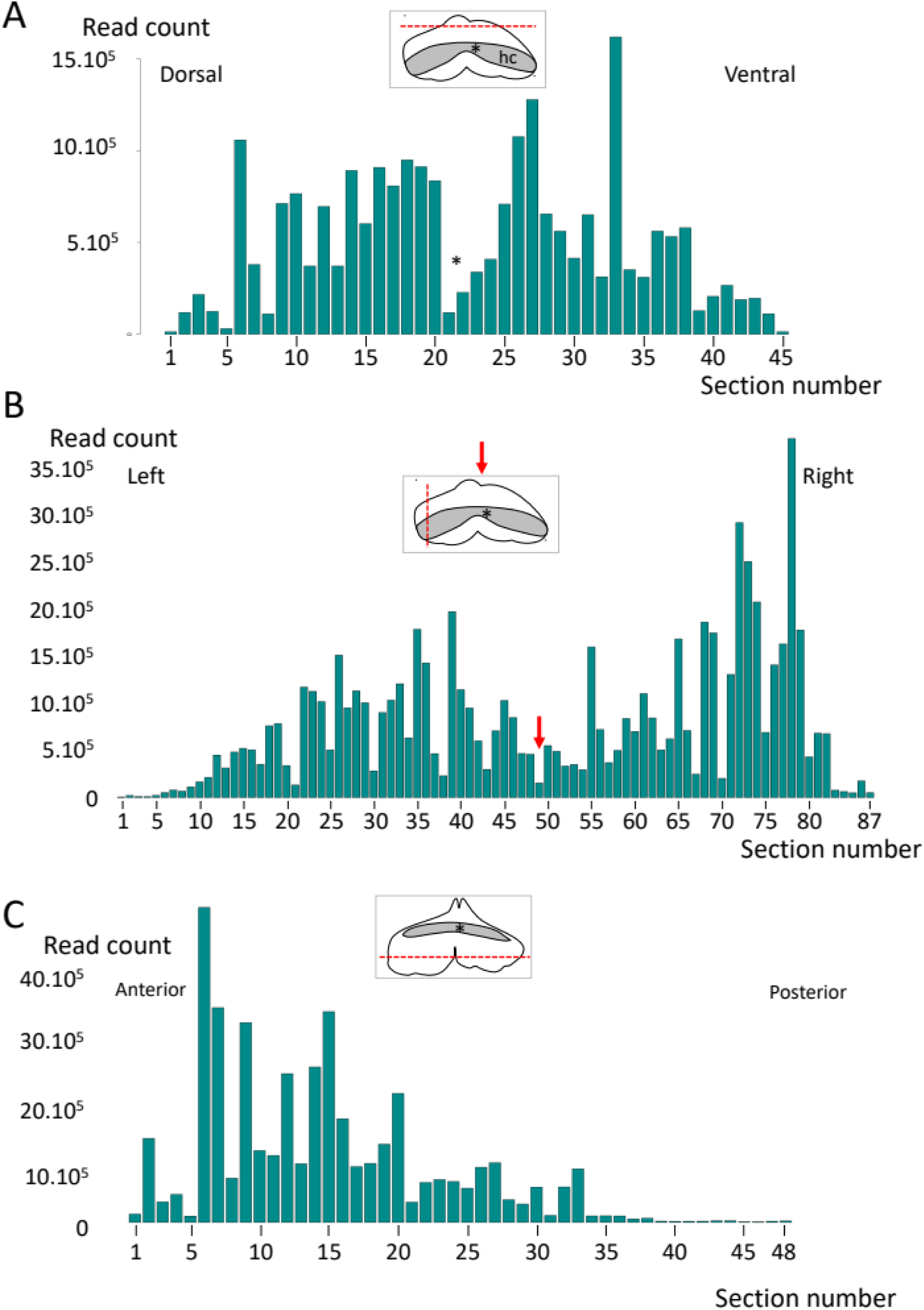
Read counts per section. **(A-C)** show read counts (y-axis) per section (x-axis) for horizontal sections (A), sagittal sections (B) and transverse sections (C). Schemes boxed in gray show front views (A,B) and a dorsal view of the habenulae, with the location of the habenular commissure (asterisk) in gray and an arrow indicating the midline. Lower reads counts are observed in the first and last sections through the organ, and the level of the HC and midline.

**Supplementary Figure 4.**
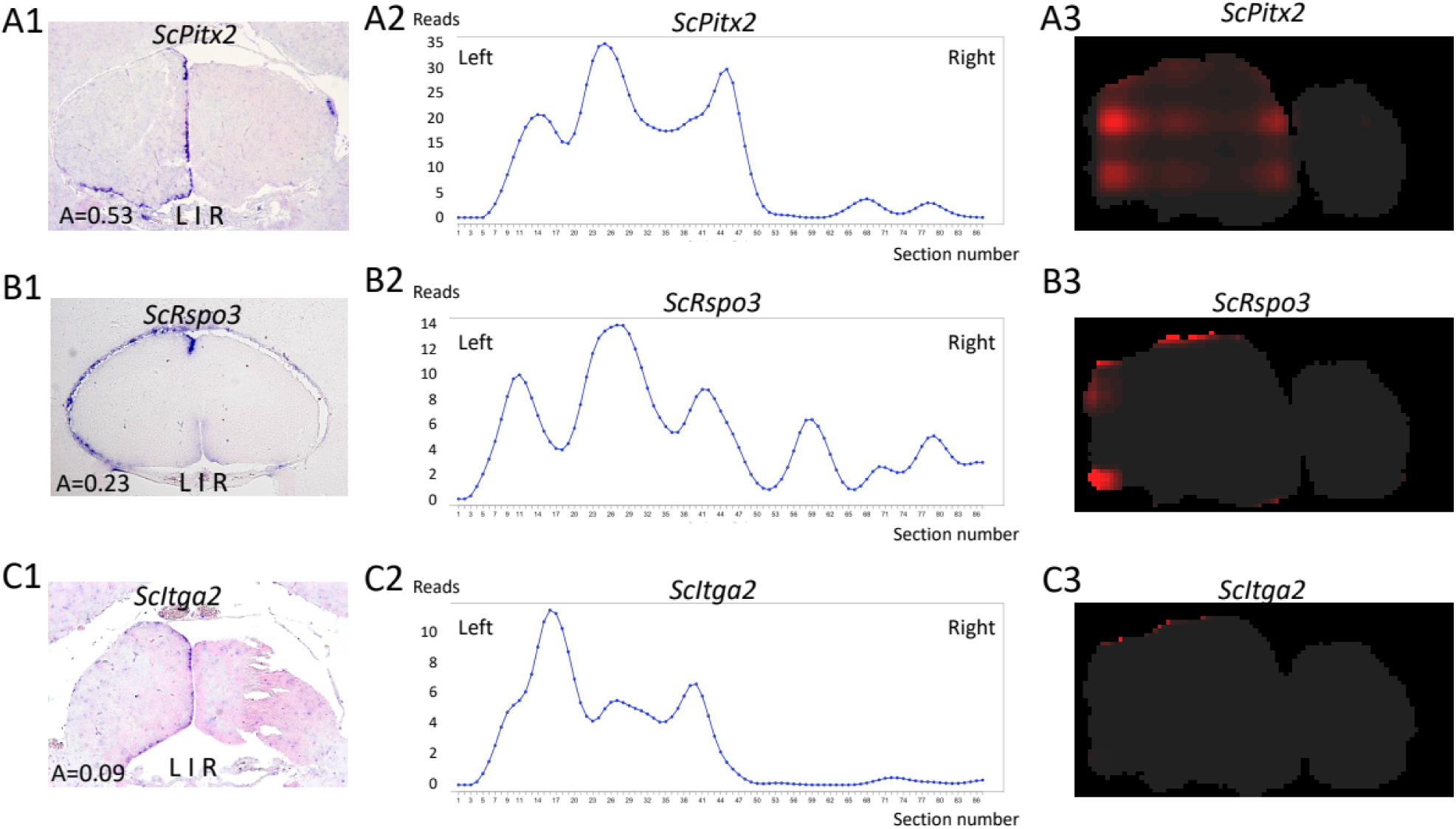
ISH and digital profiles of *ScPitx2* (A), *ScRspo3* (B), *ScItga2*. **(C)**. (A1,B1,C1) show transverse sections of catshark juvenile (A1,C1) and stage 33 (B1) habenulae following ISH. (A2,B2,C2) read counts per section along the sagittal axis and (A3,B3,C3), a digital transverse section extracted from the 3D reconstruction. These genes are expressed in the LVZ, a territory which is difficult to resolve by the iterative proportional fitting algorithm. This accounts for the inaccuracy of the 3D profiles observed, even if a higher digital signal is observed in the periphery of the habenulae for *ScRspo3* and *ScItga2*. Such patterns are not expected to yield high autocorrelation values, as observed (Cf Figure 3 and corresponding main text).

**Supplementary Figure 5.**
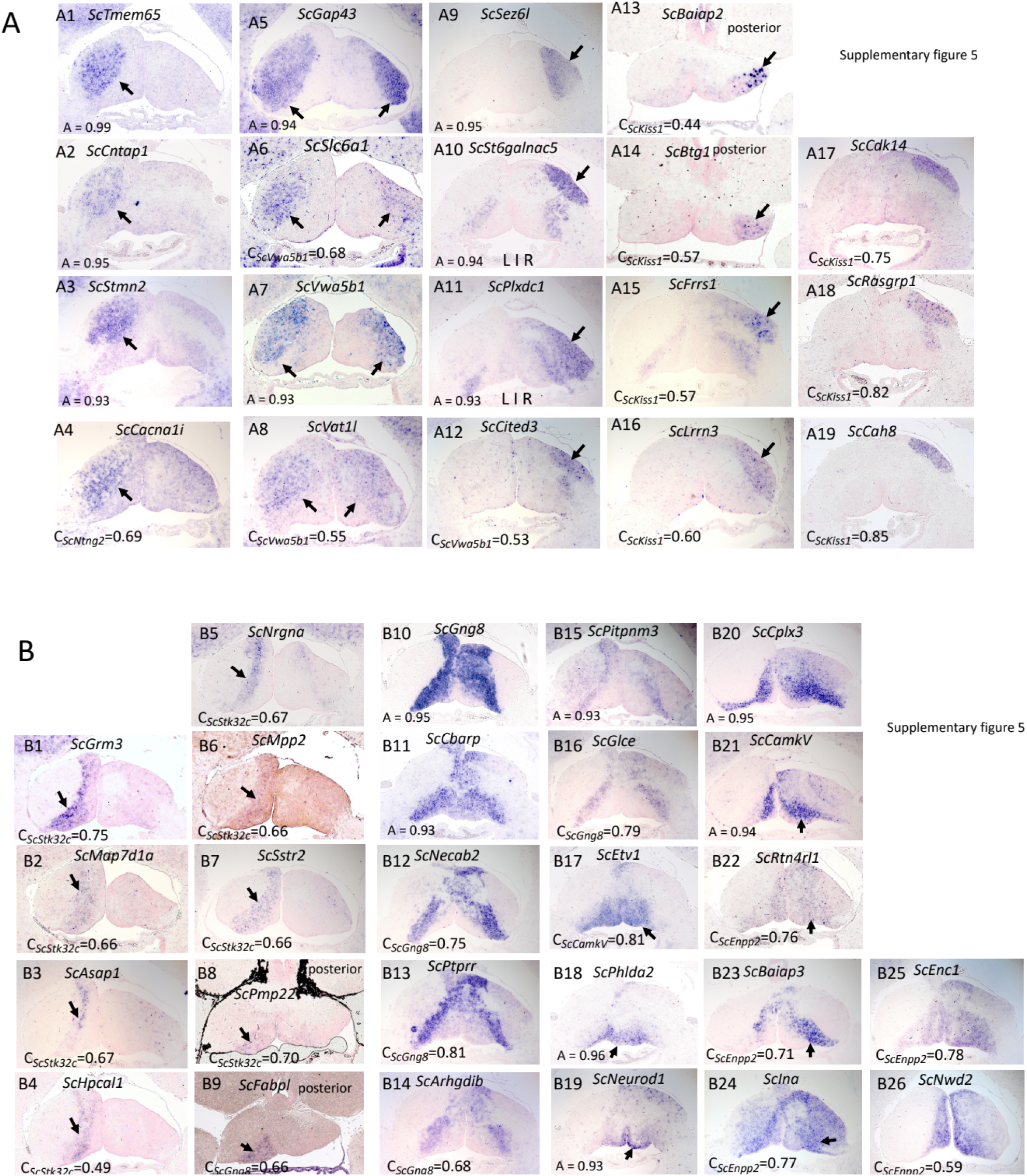
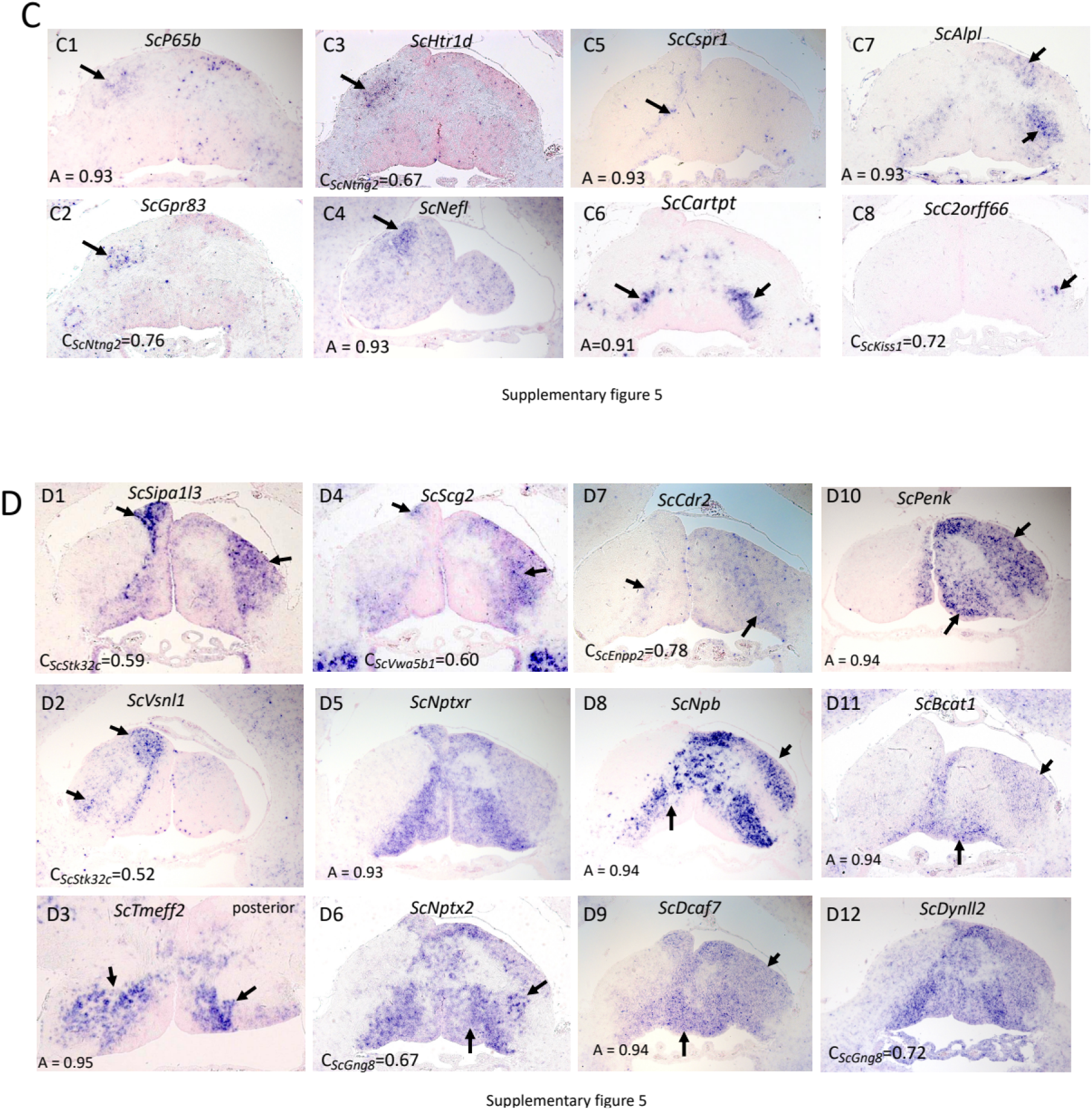
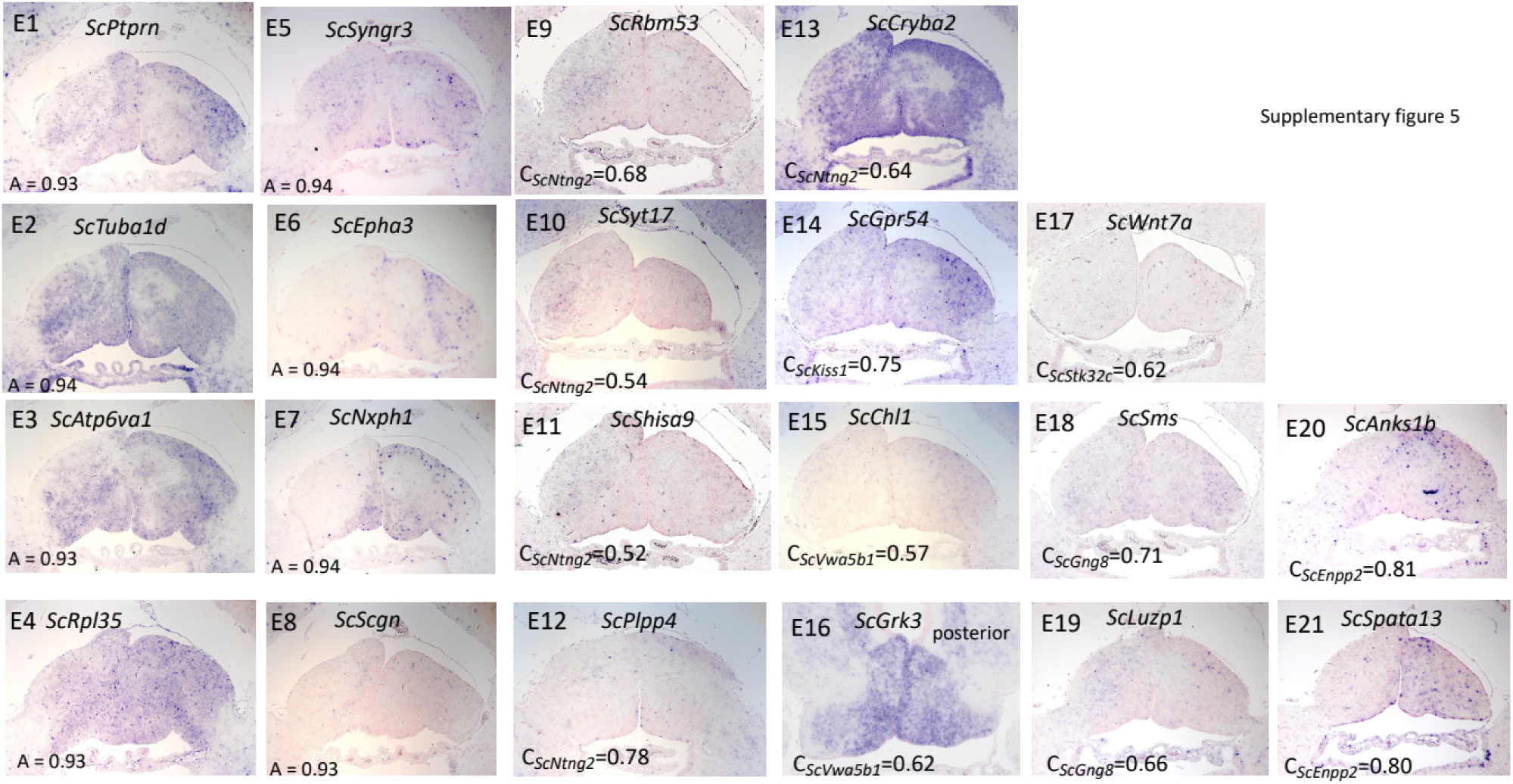
ISH expression profiles of all novel genes tested. **(A-E)** show sections of catshark juvenile habenulae after ISH with probes for the genes indicated at the top of each panel. (A) and (B) respectively show profiles spanning lateral territories (LHb, Left- and Right-LHb; A) and medial ones (MHb, MHbl, MHbm, Left-MHbl, Right-MHbl; B). (C) and (D) show profiles restricted to subdomains (C) or combinations (D) of these territories. Non-regionalized, ubiquitous or low intensity signals were also observed (E). For each gene, the criterion that was taken into account to select it for ISH analysis, either autocorrelation (A=) or correlation score with a reference gene (C_Reference_=), is indicated in the bottom left corner. All sections are transverse sections except A13, A14, B8, B9 and E16, which are horizontal sections.

**Supplementary Figure 6.**
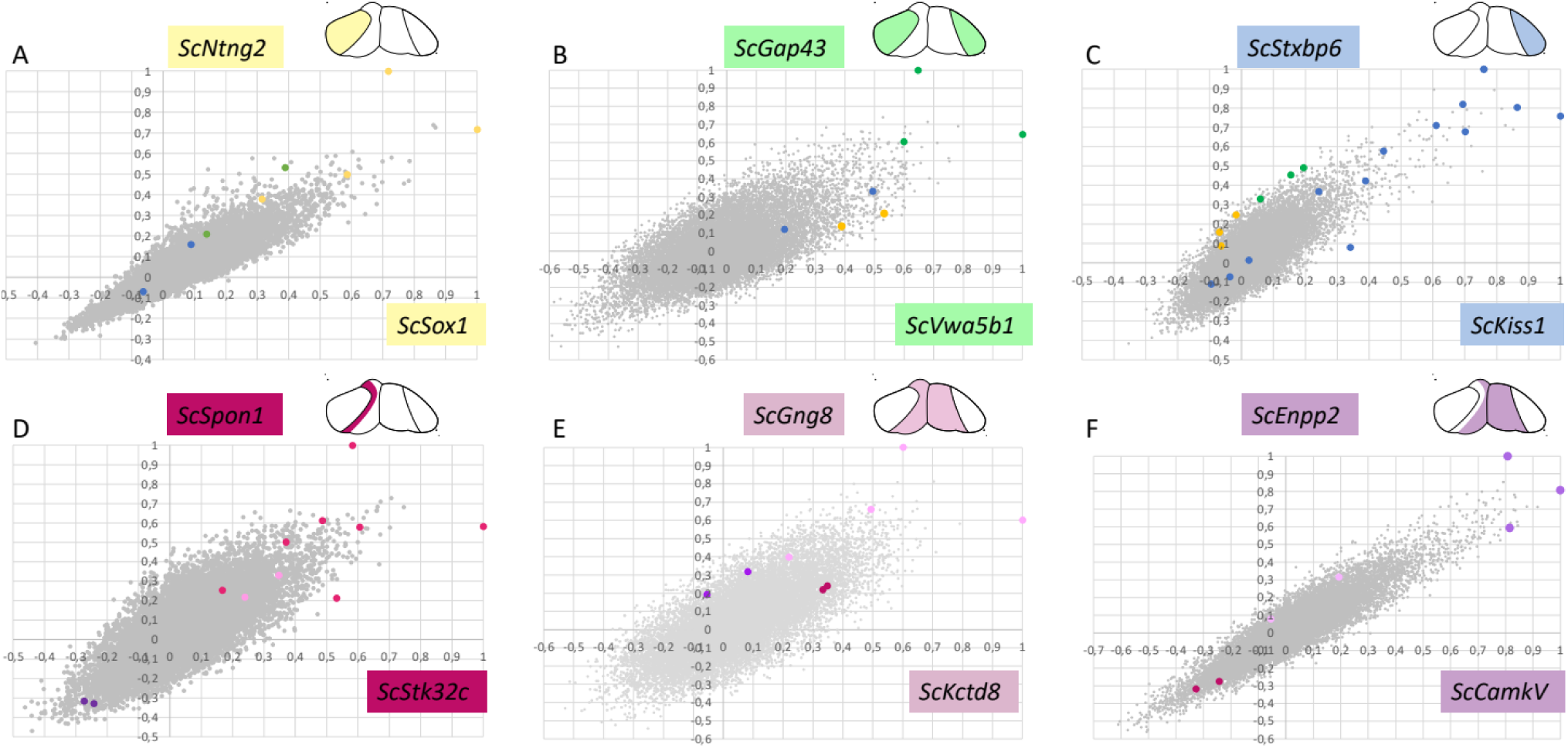
Co-variation between correlation scores for markers of the same territory. **(A-F)** Dot plots showing, for each gene model, its correlation scores (along x- and y-axes) with the following reference markers: (A), the Left-LHb markers *ScSox1* and *ScNtng2*; (B), the LHb markers *ScVwa5b1* and *ScGap43*; (C), the Right-LHb markers *ScKiss1* and *ScStxbp6*; (D), the Left-MHb markers *ScStk32c* and *ScSpon1*; (E), the MHb markers *ScKctd8* and *ScGng8;* (F), the Right-MHb markers *ScCamkV* and *ScEnpp2*. The corresponding territories are schematized at the top of each panel. Dots for known markers of habenulae are shown in colour, with the following code: Left-LHb, yellow; LHb, green; Right-LHb, blue; Left-MHb, dark magenta; MHb, pink; Right-MHb, purple.

**Supplementary Figure 7.**
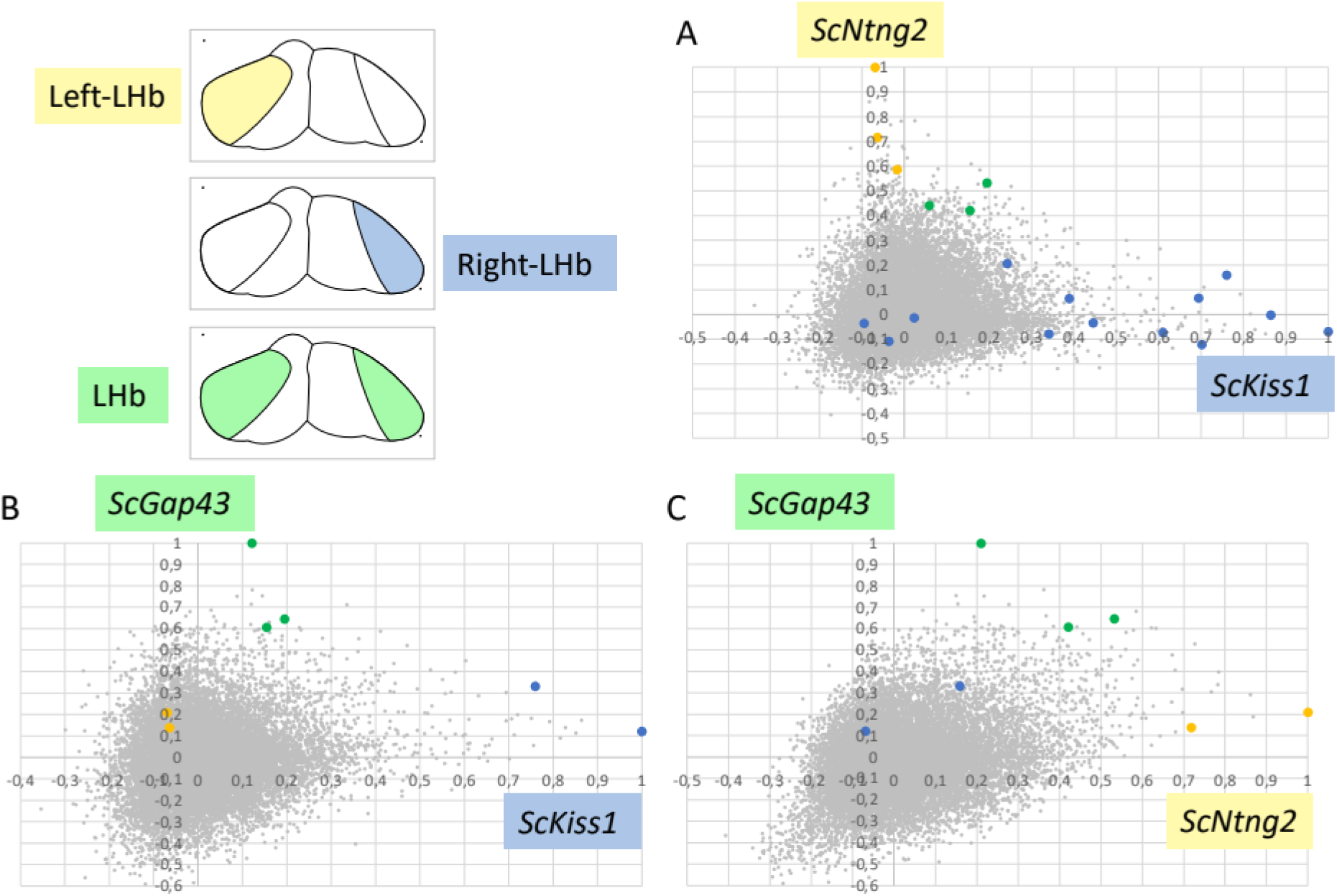
Comparison of correlation scores between markers of lateral territories. **(A-C)** Dot plots showing, for each gene model, its correlation scores (along x- and y-axes) with the following reference markers: (A), the Right-LHb marker *ScKiss1* and the Left-LHb marker *ScNtng2*; (B) the Right-LHb marker *ScKiss1* and the LHb marker *ScGap43*; (C), the Left-LHb marker *ScNtng2* and the LHb marker *ScGap43.* The territories are schematized in the top, right panel with the following color code: Left-LHb, yellow; LHb, green; Right-LHb, blue. Dots for known markers of habenulae are shown in the dot plots with the same color code. As expected, genes showing high correlations with either *ScKiss1* or *ScNtng2* tend to show low correlations with the marker of the contralateral territory. This effect is not observed to the same degree in comparisons with *ScGap43*.

**Supplementary Figure 8.**
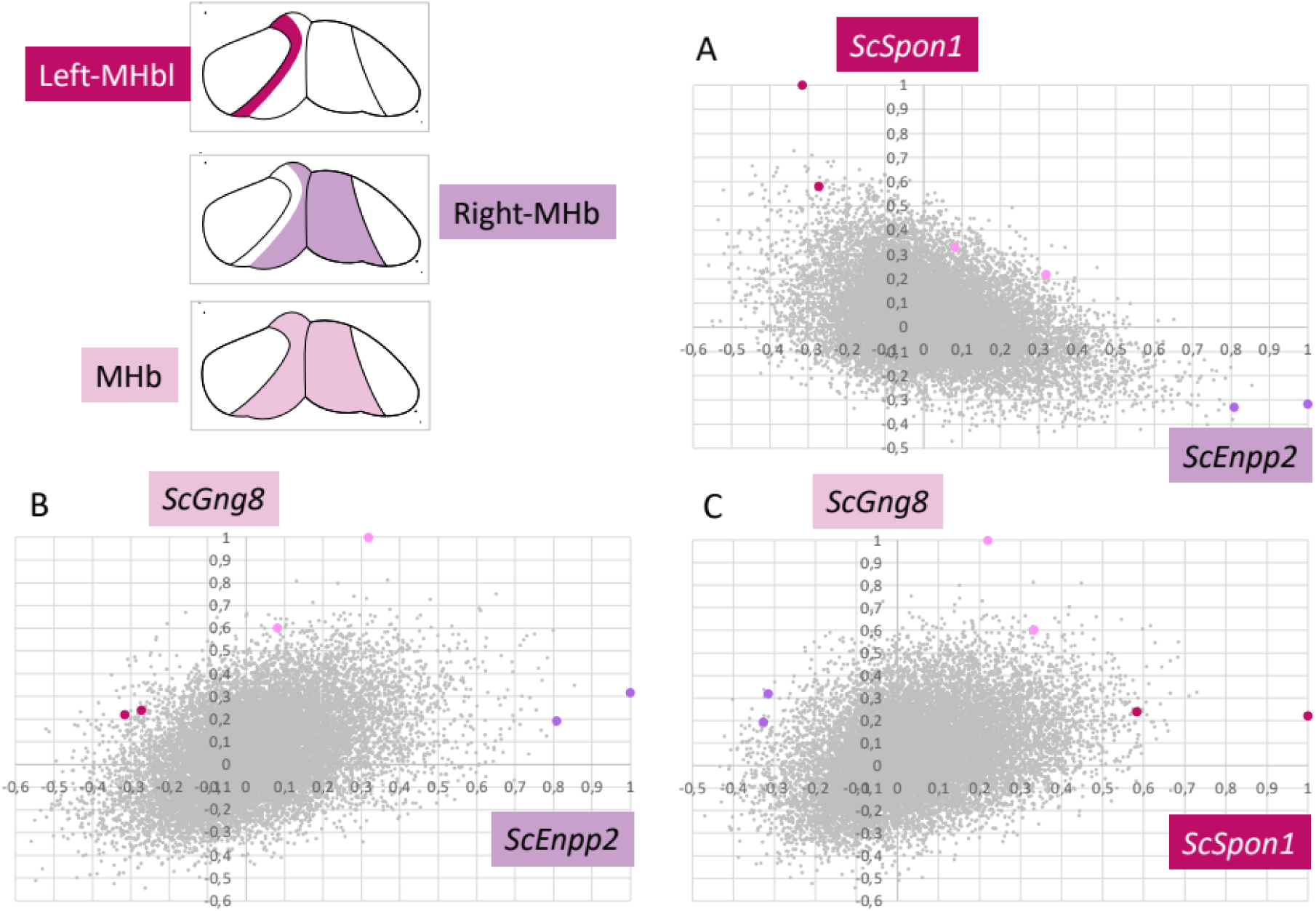
Comparison of correlation scores between markers of medial territories. **(A-C)** Dot plots showing, for each gene model, its correlation scores (along x- and y-axes) with the following reference markers: (A), the Right-MHb marker *ScEnpp2* and the Left-MHb marker *ScSpon1*. (B), the Right-MHb marker *ScEnpp2* and the MHb marker *ScGng8*. (C), the Left- MHb marker *ScSpon1* and the MHb marker *ScGng8*. The territories are schematized in the top, right panel with the following color code: Left-MHb, dark magenta; MHb, pink; Right-MHb, purple. Dots for known markers of habenulae are shown in the dot plots with the same color code. As expected, genes showing high correlations with either *ScEnpp2* or *ScSpon1* tend to show low correlations with the marker of the contralateral territory, although this effect is less marked than for lateral territories, possibly due to overlaps between digital profiles. This effect is not observed in comparisons with *ScGng8*, whose digital profile largely overlaps with those of *ScEnpp2* and *ScSpon1*.

**Supplementary Figure 9.**
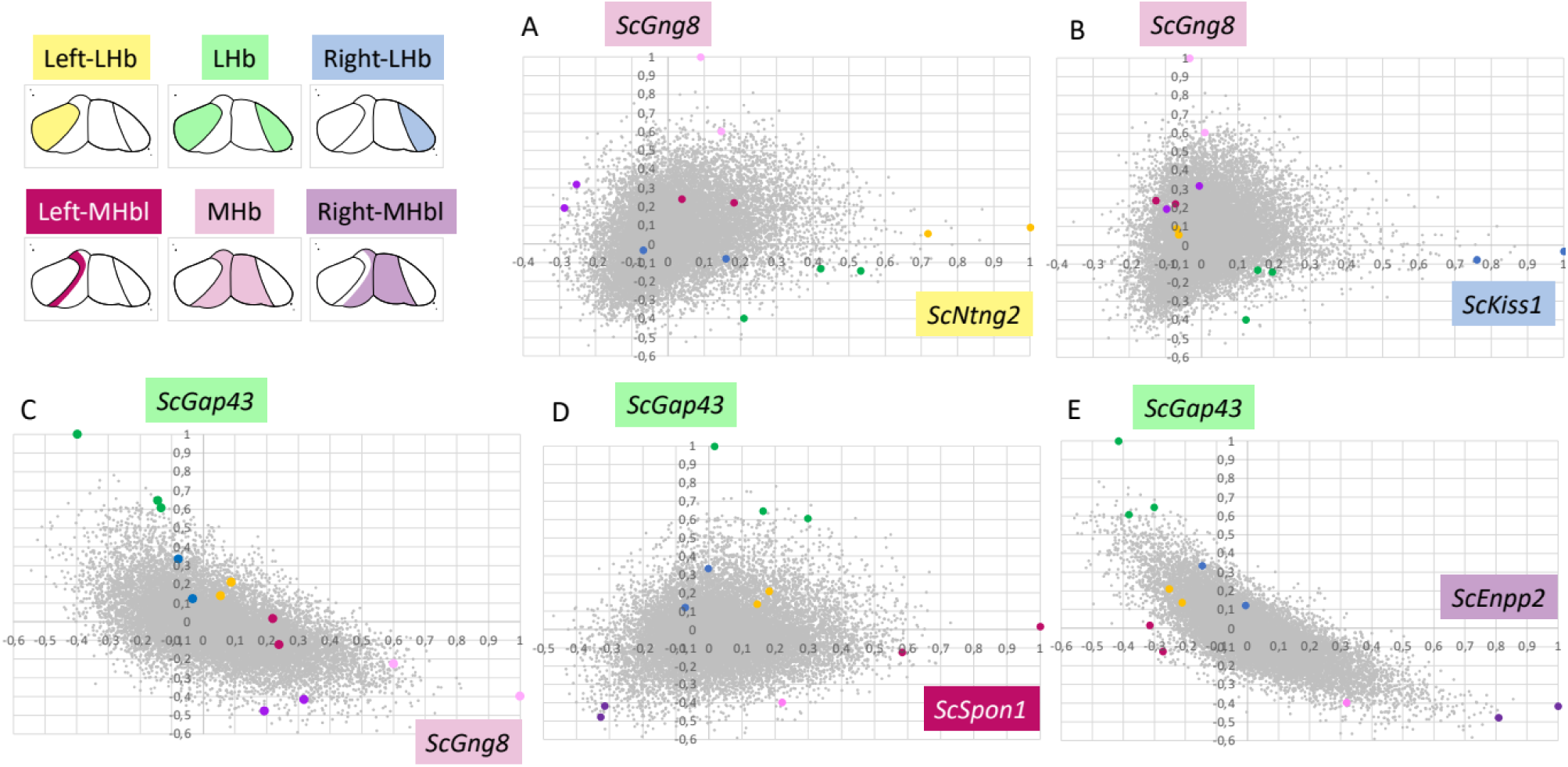
Comparison of correlation scores between markers of lateral versus medial territories. **(A-E)** Dot plots showing, for each gene model, its correlation scores (along x- and y-axes) with the following reference markers: (A), the Left-LHb marker *ScNtng2* and the MHb marker *ScGng8*; (B), the Right-LHb marker *ScKiss1* and the MHb marker *ScGng8*; (C), the LHb marker *ScGap43* and the MHb marker *ScGng8*; (D), the LHb marker *ScGap43* and the Left-MHb marker *ScSpon1*; (E), the LHb marker *ScGap43* and the Right-MHb marker *ScEnpp2*. The territories are schematized in the top, right panel with the following color code: Left-LHb, yellow; LHb, green; Right-LHb, blue; Left-MHb, dark magenta; MHb, pink; Right-MHb, purple. Dots for known markers of habenulae are shown in the dot plots with the same color code. A clear inverse variation is observed as expected between correlations with *ScGap43* and with either *ScEnpp2* or *ScGng8.* This effect is less obvious in (A,B,D) even though genes highly correlated to a lateral marker tend to have low correlations with a medial one.

**Supplementary Table 1.**
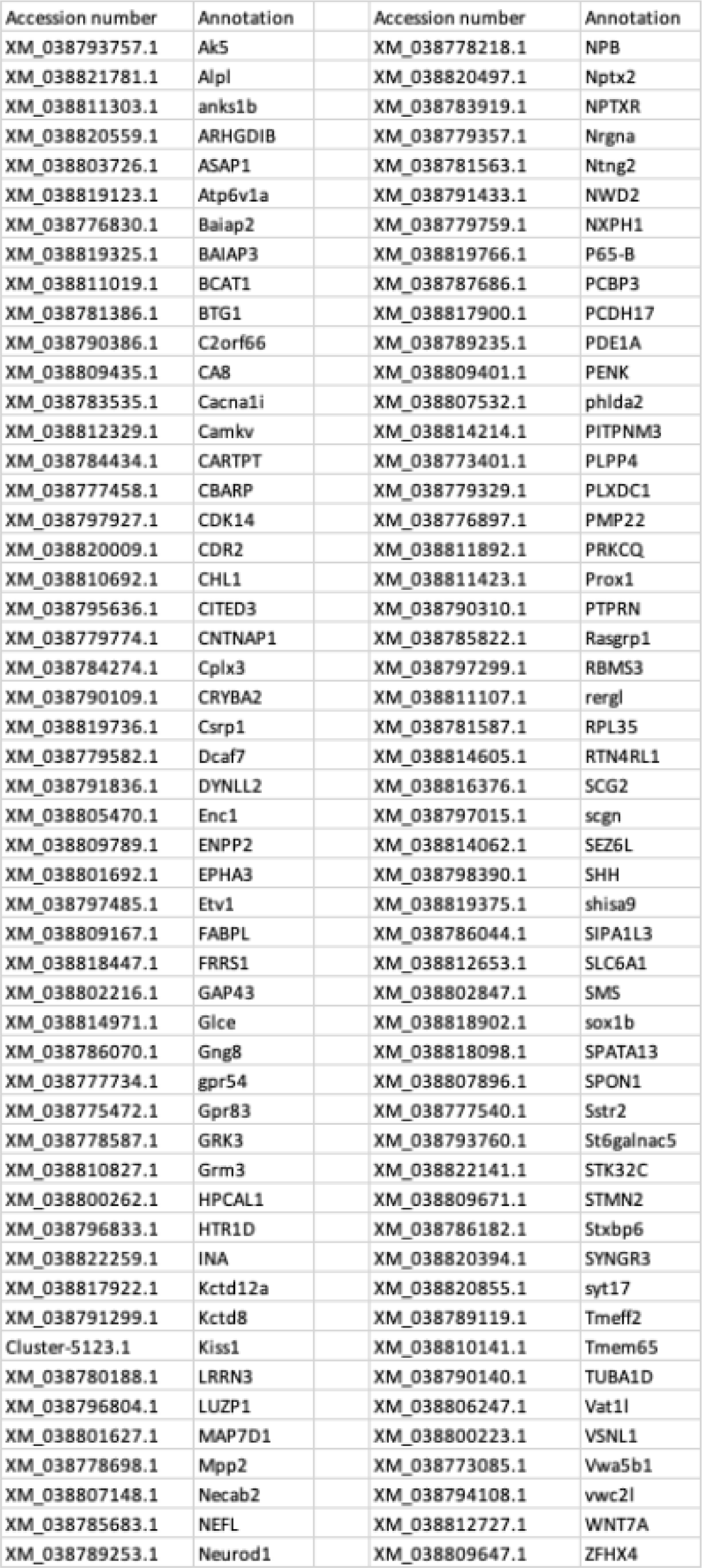
Identifiers of the gene models analyzed in this study.

**Supplementary Table 2.**
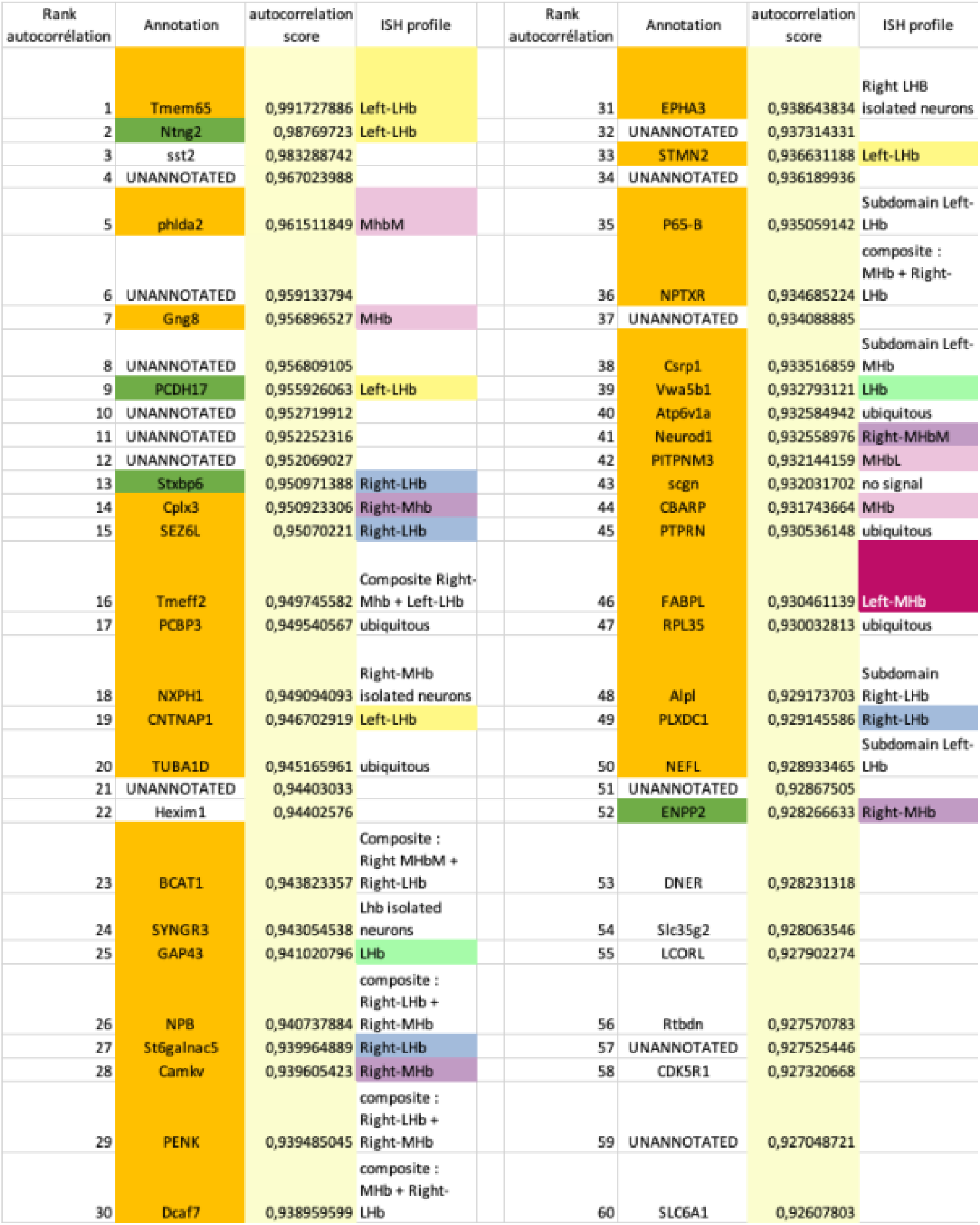
List of the 60 highest ranking genes based on autocorrelation scores. Autocorrelation rank is indicated in column A, gene name in column B, autocorrelation score in column C, and ISH profile in column D. Genes shaded in green were previously described, genes shaded in yellow were selected for ISH based on autocorrelation score. The following color code indicates expression territories as inferred from ISH: Left-LHb, light yellow; LHb, light green; Right-LHb, dark blue; Left-MHb, dark magenta; MHb, pink; Right-MHb, purple.

**Supplementary Table 3.**
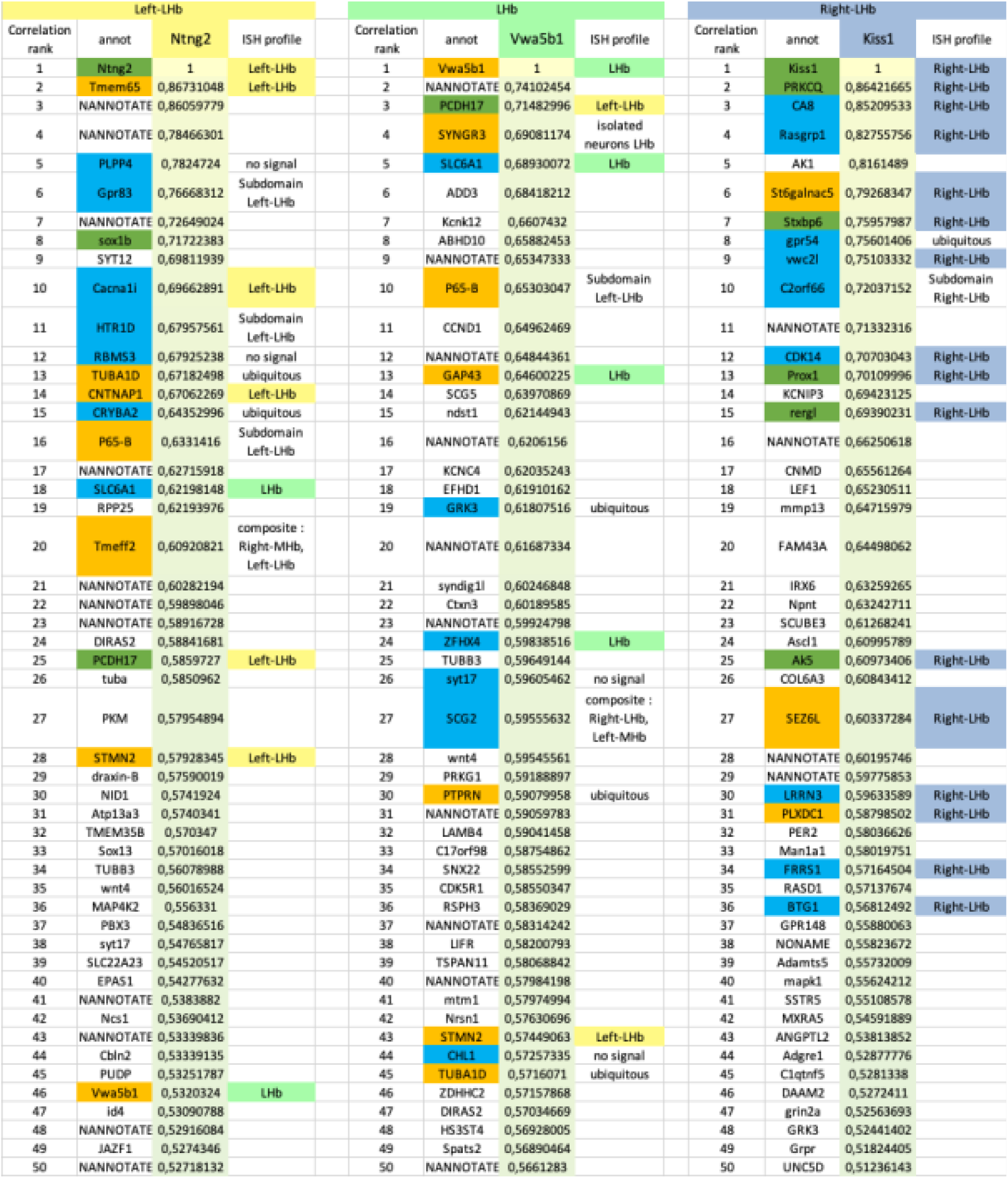
List of the 50 highest ranking genes based on correlation score with LHb markers. Correlation rank are indicated in columns A, F, K and corresponding gene name respectively in columns B, G, L. Reference gene and correlation scores are presented in columns C, H, M. ISH profiles are presented in columns D, I, N. Genes shaded in green were previously described. Genes shaded in yellow were selected for ISH based on autocorrelation score that have been tested in this study. Genes shaded in blue were selected based on correlation score with lateral reference markers. The following color code indicates expression territories as inferred from ISH: Left-LHb, light yellow; LHb, light green; Right-LHb, dark blue.

**Supplementary Table 4.**
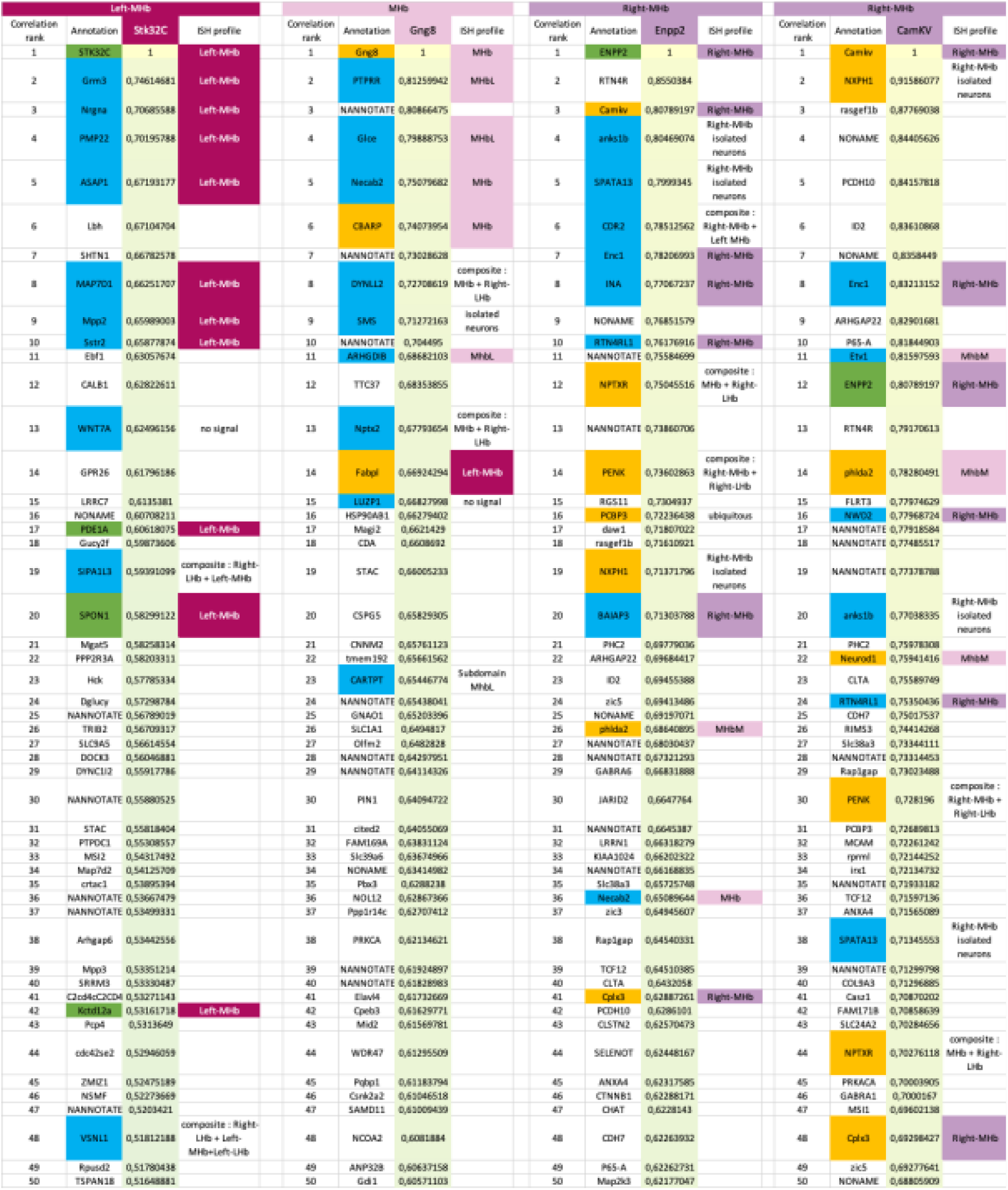
List of the 50 highest ranking genes based on correlation scores with MHb markers. Correlation rank are indicated in columns A, F, K, P and corresponding gene names respectively in columns B, G, L, Q. Reference gene and correlation scores are presented in columns C, H, M, R. ISH profiles are presented in columns D, I, N, S. Genes shaded in green were previously described. Genes shaded in yellow were selected for ISH based on autocorrelation score. Genes shaded in blue were selected based on correlation score with medial reference markers. The following color code indicates expression territories as inferred from ISH: Left-MHb, dark magenta; MHb, pink; Right-MHb, purple.

## Notes

### Competing Interest Statement

The authors have declared no competing interest.

